# A multiomics analysis of direct interkingdom dynamics between influenza A virus and *Streptococcus pneumoniae* uncovers host-independent changes to bacterial virulence fitness

**DOI:** 10.1101/2022.08.08.502690

**Authors:** Maryann P. Platt, Yi-Han Lin, Trevor Penix, Rosana Wiscovitch-Russo, Isha Vashee, Chris A. Mares, Jason W. Rosch, Yanbao Yu, Norberto Gonzalez-Juarbe

**Author notes:** Corresponding Author: Norberto Gonzalez-Juarbe, 9605 Medical Center Drive Suite 150, Rockville, MD 20850.

## Abstract

**Background:** For almost a century, it has been recognized that influenza A virus (IAV) infection can promote the development of secondary bacterial infections (SBI) mainly caused by *Streptococcus pneumoniae* (*Spn*). Recent observations have shown that IAV is able to directly bind to the surface of *Spn*. To gain a foundational understanding of how direct IAV-*Spn* interaction alters bacterial biological fitness we employed combinatorial multi-omic and molecular approaches.

**Results:** Here we show IAV significantly remodels the global transcriptome, proteome and phosphoproteome profiles of *Spn* independently of host effectors. We identified *Spn* surface proteins that interact with IAV proteins (hemagglutinin, nucleoprotein, and neuraminidase). In addition, IAV was found to directly modulate expression of *Spn* virulence determinants such as pneumococcal surface protein A, pneumolysin, and factors associated with antimicrobial resistance among many others. Metabolic pathways were significantly altered leading to changes in *Spn* growth rate. IAV was also found to drive *Spn* capsule shedding and the release of pneumococcal surface proteins. Released proteins were found to be involved in evasion of innate immune responses and actively reduced human complement hemolytic and opsonizing activity. IAV also led to phosphorylation changes in *Spn* proteins associated with metabolism and bacterial virulence. Validation of proteomic data showed significant changes in *Spn* galactose and glucose metabolism. Furthermore, supplementation with galactose rescued bacterial growth and promoted bacterial invasion, while glucose supplementation led to enhanced pneumolysin production and lung cell apoptosis.

**Conclusions:** Here we demonstrate that IAV can directly modulate *Spn* biology without the requirement of host effectors and support the notion that inter-kingdom interactions between human viruses and commensal pathobionts can promote bacterial pathogenesis and microbiome dysbiosis.

## Introduction

During annual epidemics, influenza virus causes up to 3-5 million infections and results in hundreds of thousands of deaths worldwide[1]. Disease complications and the healthcare burden rises with the development of secondary bacterial infection (SBI), which results in aggravated illness and higher mortality[2]. The major example of such synergy was observed during the 1918 Flu pandemic, in which *Streptococcus pneumoniae* (*Spn*) was recovered in ∼95% of all fatal cases. During the 2009 influenza A H1N1 pandemic, up to 43% of lung specimens recovered from fatal cases also had bacterial infection[3, 4]. SBIs are mainly caused by the Gram-positive bacterium, *Spn*[5, 6], the most common cause of community acquired pneumonia (CAP)[7]. *Spn* normally exists as a commensal in the human nasopharynx and is found in more than 95% of children under age 2, 40% of children under age 5[8] and up to 39% in the elderly[9]. Up to this point, most studies that characterize the lethal synergism between influenza A virus (IAV) and *Spn* mainly focus on the effect IAV infection has on the mammalian host. Major findings over the last decades are: 1) Factors enhancing bacterial adherence 2) Factors facilitating bacterial access to normally sterile sites, 3) Factors altering innate immune responses and 4) Synergism between viral and bacterial proteins to kill mammalian cells[5, 10–15]. During and after influenza infection, commensal bacteria can translocate from the upper to the lower respiratory tract and cause an exacerbated form of pneumonia[13, 16].

How direct physiological or pathogenic changes to bacteria are induced by direct human virus exposure in the absence of host mediated inflammatory mediators, has not been well described. *Spn* normally colonizes the nasopharynx asymptomatically. However, upon host infection with influenza virus, the bacterium can be triggered to disperse to the lungs and cause severe infection[8, 17]. Transcriptomics based studies have suggested that upon IAV infection, host antiviral inflammatory responses lead to differential expression of *Spn* genes promoting biofilm dispersal and translocation to the lungs[17]. In contrast, a recent report by Rowe *et al.* studying the early stages of disease (prior to development of severe infection), showed that IAV was able to bind directly to *Spn*, promoting adhesion, and increasing lethality in a mouse model of acute infection[18]. This phenomenon (direct binding of human virus to bacteria) has only been previously observed in studies of *Spn* interaction with respiratory syncytial virus (RSV) and suggests that respiratory viruses could directly modulate colonizing bacteria at a molecular level independently of host factors[19, 20]. However, the underlying molecular mechanisms for these phenomena have not been explored to date.

In this study, we investigate direct IAV-*Spn* interactions with emphasis on how the virus causes a switch in *Spn* physiology and virulence fitness without the requirement of host signals. Here we use Mass Spectrometry-based and molecular approaches to provide unique details of the changes occurring at the global protein level, as well as changes in phosphorylation patterns, protein secretion (secretome), growth and virulence after *Spn* exposure to IAV in a host independent manner.

## Results

### Influenza A virus modulates *Spn* growth and capsule production

Here we designed a model in which IAV is incubated with planktonic *Spn* (strain TIGR4) in liquid culture. A reduction in *Spn* growth was observed when incubated with IAV at a 1:1 ratio, with no changes in growth rate observed after *Spn* incubation with heat killed IAV (**Fig. 1A**). A similar effect was observed when *Spn* strains WU2, 6A10 and D39[21] (**Fig. S1**) or additional pathogens *Escherichia coli* and *Pseudomonas aeruginosa*[22] were co-incubated with IAV, but was not observed in the nosocomial pathogen *Serratia marcescens*[23] (**Fig. S2**). When *Spn* is grown on agar plates, two distinct colony morphologies can be observed under oblique transmitted light. These phenotypes are named transparent (low capsule expression) and opaque (high capsule expression)[24]. After incubation with IAV, we observed a dose-dependent increase in the transparent phenotype of *Spn* (**Fig. 1B**). Importantly, when biofilm grown *Spn* was challenged with IAV, a significant increase in dispersal was observed, suggesting that IAV promotes a switch from biofilm to planktonic phenotype (**Fig. 1C**). These observations suggest that *Spn* metabolism and capsule production are altered upon exposure to IAV. Of note, we tested viral stocks derived from embryonated eggs and tissue culture, and no stock-specific difference was seen in the observed phenotype (**Figure 1A**, **Fig. S3A**). We identified non-bacterial proteins present in the experimental media by label free-quantitative proteomics, which were mammalian, yeast, or influenza proteins mainly associated with tissue culture or bacterial culture media (**Fig. S3B**).

**Figure 1.**
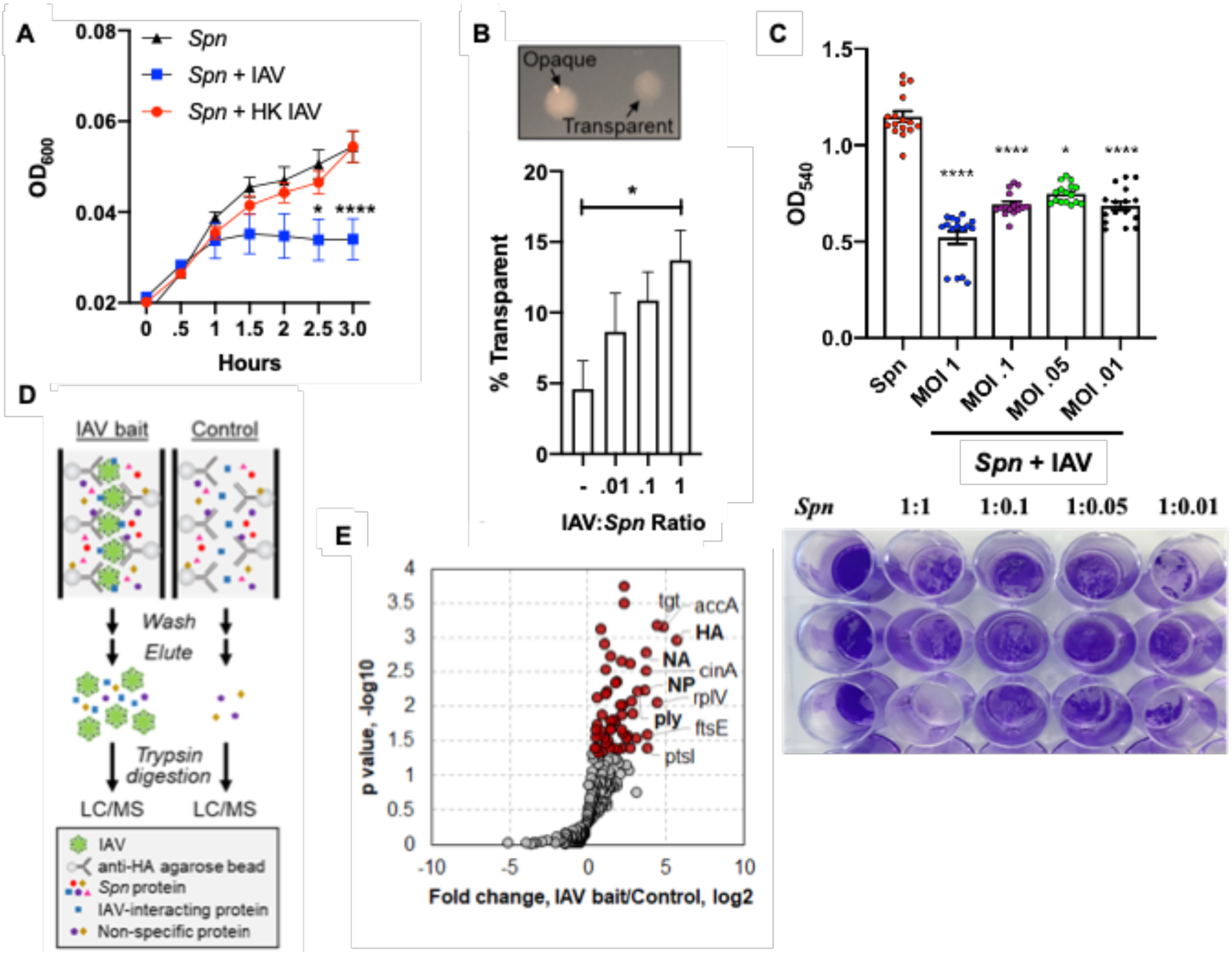
IAV-induced growth and morphological changes in *Spn*. (**A**) Growth of *Spn* strain TIGR4 in liquid media with/without IAV or with heat killed IAV (HK IAV). Data points represent mean ± SD. (**B**) Morphology of *Spn* colonies (upper panel) and counts on agar plates (lower panel) in the presence or absence of IAV. Data points represent mean ± SEM from three separate experiments. (**C**) Static biofilm of *Spn* on a polystyrene plate was dispersed upon 24h incubation with IAV. *Spn* was stained by crystal violet. Quantitation was done by measure of absorbance at 540nm. (**D**) Schematic illustration of the experimental process for Affinity Enrichment Mass Spectrometry (AE-MS). Influenza virus particles were bound to HA antibody immobilized on agarose beads, which serve as bait to enrich *Spn* proteins. The control experiment did not contain the IAV bait. (**E**) Volcano plot showing proteins identified in the AE-MS analysis. Proteins significantly enriched in the presence of IAV bait (Student’s t-test, p<0.05) are shown in red. IAV proteins are in bold. Kruskal-Wallis test with Dunn’s multiple-comparison post-test. Asterisks denote the level of significance observed: * = p ≤ 0.05; ** = p ≤ 0.01; *** = p ≤ 0.001.

### Proteins as mediators of IAV-*Spn* interaction

Previous work by Rowe *et al.* showed direct interaction between *Spn* and IAV independent of both the glycosylation status of the virus and sialyation on the surface of *Spn*[18]. These data suggest that surface proteins could mediate the observed *Spn*-IAV interaction. Therefore, we used the Affinity Enrichment Mass Spectrometry (AE-MS) approach[25]to search for putative IAV-interacting *Spn* proteins, (**Fig. 1D**). IAV particles mixed with anti-Hemagglutinin (HA) agarose beads were incubated with *Spn* cell lysate. *Spn* proteins that can bind to the virus particles are retained on the beads. After extensive washes, proteins were eluted and analyzed by LC-MS. To exclude *Spn* proteins that bind non-specifically to the beads and resulted in background identification, we performed a control experiment without the addition of IAV particles to the antibody coated beads. Each experiment was performed in triplicate and protein quantitation was performed using the label-free quantitation (LFQ) approach.

Both the control and experimental group comprise of 200-300 quantifiable proteins, suggesting high background from non-specific interactions of *Spn* proteins to anti-HA beads. IAV proteins, Hemagglutinin (HA), Neuraminidase (NA), and Nucleoprotein (NP), were confidently identified in the experimental group but not in the control group, indicating successful coating of anti-HA beads with IAV particles. After statistical analysis (Student’s t-test, p < 0.05), 53 *Spn* proteins were found to be significantly enriched in the experimental group (i.e., with IAV particles) (**Fig. 1E, Spreadsheet S1**), 42 of which were increased by more than 2-fold. Of interest, one of the enriched proteins was pneumolysin (Ply) (increased by nearly 3-fold, **Fig. 1E**). Ply is a cholesterol-dependent pore-forming toxin that forms lytic pores on target membranes and modulates host inflammatory responses during *Spn* infection[26, 27]. Ply is known to be released by *Spn* upon autolysis and associate with host membranes via interaction with cholesterol[28]. Here, Ply from the cytosol may have been released upon *Spn* lysis to bind to IAV. However, a previous study indicated that Ply can localize to the *Spn* cell wall in a non-autolytic way, providing a mechanism for meaningful *Spn*-IAV interaction via Ply[29]. Other proteins found to bind influenza, and that are found in *Spn*’s cell or cell surface accessible include gapN, ptsl, accA, ftsE and cinA, among others (**Fig. 1E, Spreadsheet S1**). Our AE-MS approach identified candidate *Spn* proteins mediating the *Spn*-IAV interaction, and the downstream molecular mechanisms of the interactions between proteins need to be further studied in detail.

### Proteome profile of *S. pneumoniae* is directly altered by influenza A virus

To gain a better understanding of the molecular changes to *Spn* upon direct interaction with IAV, we profiled the global *Spn* proteome with and without co-culture with IAV for 1 hour at a 1:1 ratio. The cell pellets (each in biological triplicates) were processed following the Suspension Trapping (STrap) approach with in-house assembled filter devices[30]. The resulting peptides were subjected to direct liquid chromatography–tandem mass spectrometry (LC-MS/MS) analysis for global proteomics, or titanium dioxide (TiO2) based phosphopeptide enrichment and then LC-MS/MS analysis for phosphoproteomics (**Fig. 2A**). The MaxLFQ based label-free quantitation (LFQ) was performed to determine proteome-level changes in response to IAV challenge[31]. Overall, 937 *Spn* proteins were quantified (≤ 1% FDR on both protein and peptide level), comparable to the quantifiable *Spn* proteome size of 919 proteins reported by a recent study[32]. In another independent set of experiments, we identified 1,036 quantifiable proteins, with 85% of them commonly quantified, indicating good experimental reproducibility (**Fig. S4A**). The overall proteome spanned around five orders of magnitude (**Fig. S4B**). Correlation between the biological replicates was high (Pearson ***r*** = 0.97 ± 0.01; n = 6. **Fig. S3C**), whereas the inter-group correlation was low (0.75 ± 0.02; n = 9). The data implied that the IAV challenge-dependent proteome-level alternations could be captured by our approach, thereby suggesting its general applicability for quantitative proteomics. Of the quantifiable *Spn* proteins (in at least 2 of 3 replicates), 378 of them showed a significant difference between the two groups (Student’s t-test, p < 0.05), including 104 and 274 proteins up- and down-regulated (fold change ≥ 1.5) in the presence of IAV, respectively (**Fig. 2B**, **Spreadsheet S2**). Their differential expressions were also verified in the second set of experiments (**Fig. S4D**). This finding provides the first global view of how IAV alters the *Spn* proteome, showing that one-third of quantifiable proteins had altered expression. Kyoto Encyclopedia of Genes and Genomes (KEGG) pathway enrichment analysis showed that approximately one third of the significant proteins are involved in metabolic pathways (FDR = 9.30E-13) (**Fig. 2C**), suggesting IAV directly influences *Spn* metabolism. Proteins associated with carbohydrate, amino acid, and nucleotide metabolism pathways were downregulated in the presence of IAV (**Fig. 2C**), consistent with the slower growth phenotype (**Fig. 1A**). Purine and pyrimidine synthesis are central to many metabolic processes of *Spn*[33], including DNA synthesis and capsule production[34, 35]. Our proteomic analysis showed 9 of the purine biosynthesis enzymes (PurB, PurC, PurD, PurE, PurF, PurH, PurK, PurM, and PurN) and 11 of the pyrimidine biosynthesis proteins (CarA, CarB, PyrB, PyrC, PyrDA, PyrDB, PyrE, PyrF, PyrH, PyrK, and PyrR) were down-regulated upon IAV challenge (**Fig. S5 and Fig. S6**), many of them by more than 10-fold. These observations directly correlated to early transcriptional changes observed after incubation of *Spn* with IAV (**Fig. 2D**, **Spreadsheet S3**). Moreover, proteins associated with amino acid biosynthesis pathways (**Fig. S7**) and transport of amino acids (e.g., AliA, AliB, AmiE, AmiF) were also downregulated in the presence of IAV. Among the significantly down-regulated proteins, we also found several capsule synthesis proteins, including CpsE, CpsF, CpsG, and CpsK[36] (**Fig. S8**), indicating reduced capsule production and consistent with our observation that more transparent colonies were present on agar plates after *Spn* co-culture with IAV (**Fig. 1B**). In contrast, proteins that control cell division, including FtsA, ABC transporters FtsE, FtsX, and DivIB[37–39], were found up-regulated (**Fig. S8**).

**Figure 2.**
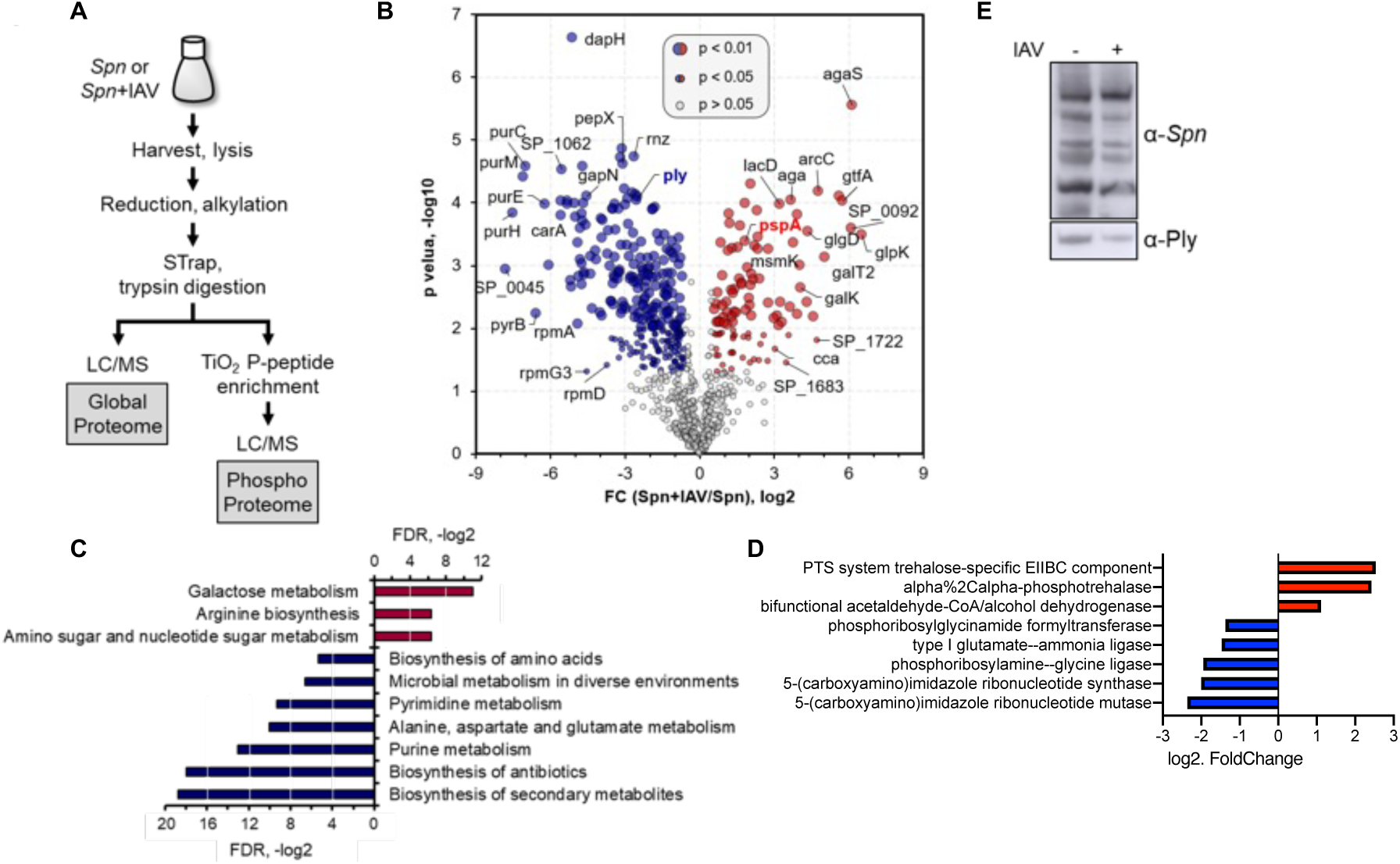
Global proteomic change of *Spn* in the presence of Influenza virus. (**A**) Workflow for global and phosphoproteome profiling of *Spn*. (**B**) *Spn* proteins identified from the global proteome analysis shown in a volcano plot. Red and blue are proteins significantly up- or down- regulated by 1.5-fold when co-incubated with IAV (Student’s t-test, p<0.05 or p<0.01 for sizes indicated). KEGG enrichment analysis of significant (**C**) proteins and (**D**) transcripts. (**E**) Western blot of *Spn* lysate grown in presence or absence of IAV, probed with a polyclonal anti-*Spn* or anti- Pneumolysin antibody.

Regulation of carbohydrate metabolism has been indicated to directly coordinate pneumococcal colonization and virulence[40]. We found that catabolite control protein A (CcpA), a regulator for carbohydrate utilization of *Spn*[41–43], was slightly up-regulated (1.8-fold) in the presence of IAV (**Fig. S8**), but glycolysis, the pathway *Spn* utilizes for glucose metabolism, was mainly down-regulated (**Fig. S9**). Importantly, galactose metabolism, which occurs preferentially in *Spn* that colonizes the respiratory tract[40] and cardiac tissues[44, 45] to modulate virulence, was found to be enriched among up-regulated proteins (**Fig. 2C**, **Fig. S10**). Enzymes involved in galactose metabolism pathways, the Leloir pathway (e.g. Aga, GalK, GalT2) and the tagatose 6- phosphate pathway (e.g. LacA, LacB, LacC, LacD)[46, 47], were found to be up-regulated. The complete list of proteins significantly altered in the presence of IAV can be found in **Spreadsheet S2.** Together, the change in utilization of sugars upon IAV challenge indicates not only altered metabolism, but also a potential virulence switch in *Spn*.

Supporting our virulence switch hypothesis, several virulence proteins were also found to be differentially expressed when *Spn* was challenged with IAV. The pneumococcal surface protein A (PspA), a surface-exposed virulence molecule of *Spn* that helps subvert the host innate immune response by interfering with the complement system[26, 48], was found to be almost 2-fold up- regulated in the presence of IAV (**Fig. 2B**). In contrast, Ply was found to be more than 5-fold down-regulated (**Fig. 2B**) in the presence of IAV. The reduction in Ply was further validated by immunoblot (**Fig. 2E**). Of note, of ten common proteins associated with antimicrobial resistance[49], five were found to significantly change upon IAV incubation. Downregulated factors were murD, murG, metG and rpoB, whereas murI was observed upregulated. While expression of the serine/threonine kinase StkP did not change (**Fig. S5**) its phosphorylation was significantly changed in multiple sites (**Fig. 3**), suggesting its enhanced activity. The intricate regulation of virulence factor expression upon IAV co-incubation reflects how *Spn* virulence can be modulated. Despite the change of several pathways and *Spn* virulence determinants, IAV challenge did not change expression of the energy-coupling factor transporter ATP-binding protein EcfA2 (also known as SP_2220 or CbiO2), a putative cobalt transporter which when absent has been reported to be involved in enhanced pneumococcal pathogenicity in IAV-infected animals[50]. Finally, even in the presence of IAV, there was very low expression (quantified by ≤ 3 peptides) of the oxalate/formate antiporter SP_1587, which has been shown to regulate bacteremia, neutrophil infiltration, and pulmonary damage in IAV-infected mice[51]. This might reflect a need for interaction with the host (in addition to viral proteins) to alter expression of other virulence factors and colonize the lungs without induction of excess inflammation. In summary, our global proteomic analyses substantiate our initial observations that *Spn* growth, metabolism, and virulence, are directly affected upon exposure to IAV.

**Figure 3.**
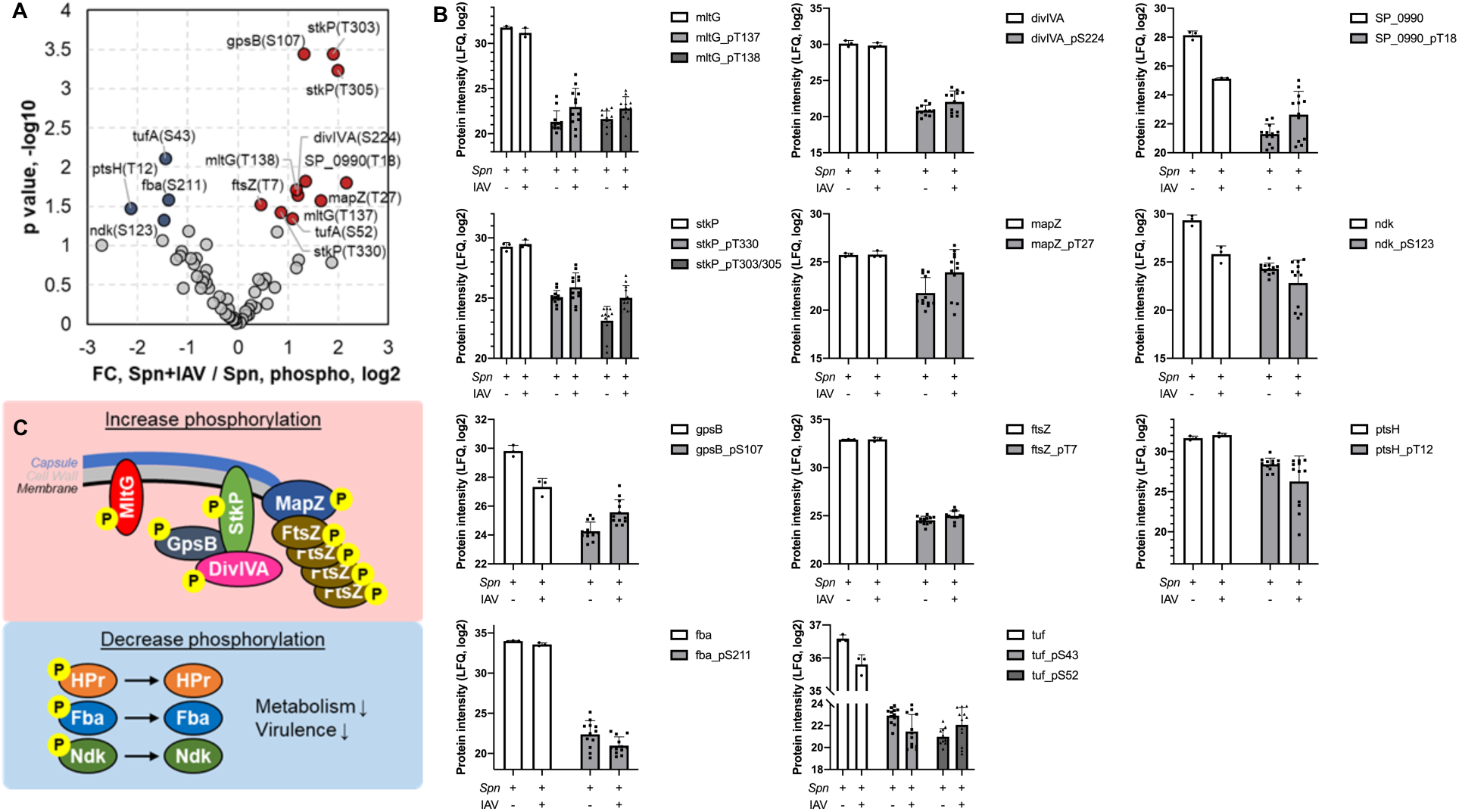
Change in *Spn* phosphoproteome in the presence of Influenza virus. (**A**) Volcano plot showing fold changes of phosphopeptides vs their p-values. Significantly changed phosphopeptides are shown in red (up-regulated) and blue (down-regulated) in the presence of IAV (Student’s t-test p<0.05). (**B**) Peptide counts of the unmodified (white) and phosphorylated (gray) forms of indicated proteins in the absence or presence of IAV. Data reflect mean ± SD. (**C**) The top scheme shows the proteins with increased phosphorylation, most of which are known to localize to the septum during cell division. The bottom blue square shows proteins that are downregulated in phosphorylation, and their related functions.

### IAV directly alters *Spn* protein phosphorylation

Phosphorylation is an important post-translational protein modification that regulates many cellular processes across all kingdoms of life[52]. In bacteria, two-component systems mediate cellular responses to environmental cues via histidine phosphorylation, while serine/threonine and tyrosine kinases also exist to modulate various cellular functions[53, 54]. In *Spn*, there is one serine/threonine kinase, StkP, and one tyrosine kinase, CpsD, identified as regulators of cell wall synthesis, cell division, and capsule synthesis[53, 55]. To further elucidate the molecular and functional changes in *Spn* upon its direct interaction with IAV, we analyzed the phosphoproteome of *Spn* after IAV challenge using a TiO2- based phosphopeptide enrichment method[56, 57] (**Fig. 2A**). Collectively, we quantified 108 class-1 phosphopeptides (probability score > 0.75)[58] derived from 55 *Spn* proteins, similar to the size of a previously reported *Spn* phosphoproteome[59]. We further required that quantifiable phosphopeptides appear in at least two out of three replicates of each group, resulting in 66 phosphopeptides from 29 *Spn* proteins (**Fig. 3A, Spreadsheet S4**). Among them, 16 were serine phosphorylation, 33 were threonine, and 17 were tyrosine phosphorylation. Both StkP and CpsD were identified in our phosphoproteome analysis with multiple phosphorylation sites. For StkP, 5 threonine residues located in the juxta-membrane region (Thr293, Thr295, Thr303, Thr305, Thr330)[53, 60] were identified, whereas in CpsD, 4 tyrosine residues (Tyr215, Tyr218, Tyr221, Tyr224) and 1 serine residue (Ser220) located in the C-terminal Y-rich domain[55, 61] were identified. Proteins that were previously found to be substrates of StkP, such as DivIVA, MapZ, and FtsZ[53, 62–64], were identified with multiple phosphorylation sites (**Table S1, Spreadsheet S4**). CpsE, a component within the capsular polysaccharide synthesis complex[55], and SP_1368, a membrane protein that contains a LytR_cpsA_psr domain, were identified with tyrosine phosphorylation, and therefore could be possible substrates of CpsD. Some phosphoproteins we identified are involved in metabolic pathways, such as Fba, Gap, GpmA (glycolysis), Ndk and NrdE (purine synthesis) (**Fig. 3A**). Additionally, other identified *Spn* phosphoproteins have never been reported before (**Table S1**). Identified phosphoproteins are involved in cell growth, capsule synthesis, and metabolism[59, 64] and were differentially modified upon *Spn* interaction with IAV.

### Further analysis of the *Spn* phosphoproteome after challenge with IAV

15 phosphopeptides were found to be significantly different between the two groups (Student’s t-test, p<0.05, **Fig. 3A**).

StkP and several of its substrates, including MapZ, DivIVA, MltG, GpsB, and FtsZ, were found to be upregulated in phosphorylation when IAV was present (**Fig. 3A, B, Table S1**). MapZ directs the FtsZ (Z-ring) assembly and constriction during cell division, while StkP arrives in the mid-cell later where it phosphorylates DivIVA and decreases peripheral peptidoglycan synthesis[53, 62, 65]. GpsB mediates localization of StkP to the septum and is required for its kinase activity[62, 65, 66] (**Fig. 3C**). The abundance level of these proteins was not changed (**Fig. 3B, Spreadsheet S2**) except GpsB, which was found to be downregulated, indicating a true increase in phosphorylation of these proteins. The function of phosphorylation at specific sites in these proteins needs to be studied further. Proteins that were found to be downregulated in phosphorylation were mostly metabolic proteins (Fba, Ndk, PtsH) (**Fig. 3B**). Notably, the phosphocarrier protein Hpr (PtsH), had the largest decrease in phosphorylation, at Thr12, by more than 4-fold. The phosphorylation of Hpr has been shown to mediate CcpA interaction with genes that regulate carbohydrate usage and virulence factor expression[41, 67]. Taken together, this is consistent with our observations in the global proteome and suggests a switch in metabolism and virulence when *Spn* is challenged with IAV (**Fig. 3C**).

### The release of *Spn* surface proteins is increased upon challenge with influenza A virus promoting evasion of complement activity

Like many other bacteria, *Spn* has many surface proteins important for colonization, virulence, and host invasion[68, 69]. Some of these surface proteins bind to cell wall phosphorylcholine, some are lipoproteins that associate with membrane transporters, and some have unknown anchoring mechanisms. The observed reduction in *Spn* capsule production after challenge with IAV (**Fig. 1B**) may lead to the release of surface proteins. To address this hypothesis, we performed proteomic profiling of the conditioned media used to grow *Spn* which contains bacterial proteins released from the cell surface upon challenge with IAV (**Fig. 4A**). Collectively, 30 *Spn* proteins were identified in the conditioned media, most of which had higher abundance (by peptide-spectrum match (PSM) count) after IAV challenge (**Spreadsheet S5**). Consistent with our hypothesis, this result suggests that IAV can trigger *Spn* surface protein release, possibly due to reduced capsulation (**Fig. 1B**).

**Figure 4.**
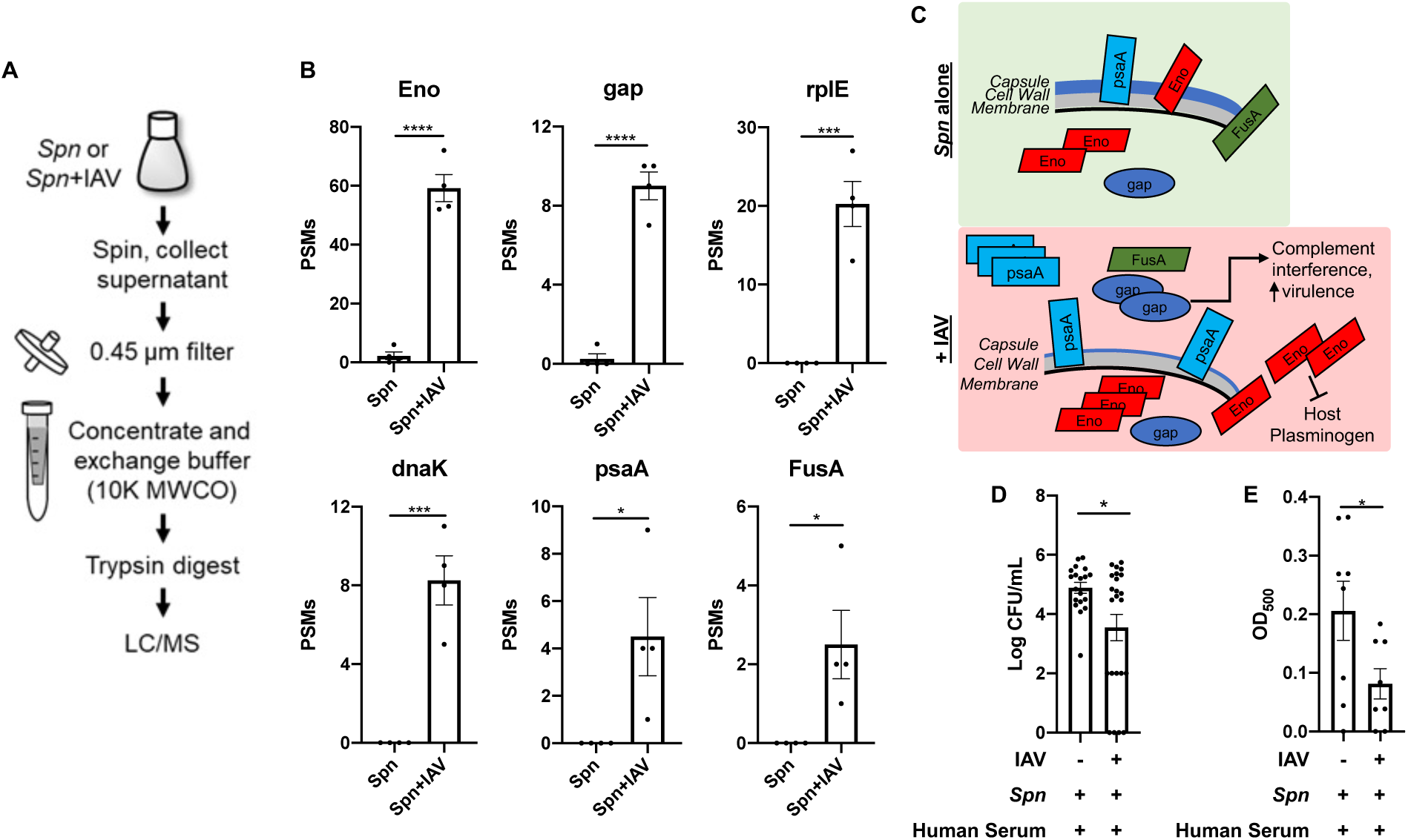
Release of *Spn* surface proteins was increased in the presence of Influenza virus and promotes complement inhibition. (**A**) Workflow for the collection and processing of extracellular *Spn* proteins for LC/MS analysis. (**B**) PSM count of extracellular *Spn* proteins in the presence or absence of IAV (Data reflect n = 4 presented as mean ± SD, Two-tailed Student’s t- test, *p <0.05, ***p<0.001, ****p<0.0001). Eno, enolase; RplE, 50S ribosomal protein L5; GAP, Glyceraldehyde-3-phosphate dehydrogenase; dnaK, Chaperone protein; PsaA, Manganese ABC transporter substrate-binding lipoprotein; FusA, Elongation factor G. (**C**) Schematic model of the proposed IAV-induced *Spn* pathway changes that lead to enhanced host invasion and inhibition of complement. (**D**) Recovered *Spn* (CFU/ml) in phagocytosis assay. (**E**) Complement-driven hemolysis assay in red blood cells challenged with *Spn* Δply. Student t test, asterisks denote the level of significance observed: * = p ≤ 0.05; ** = p ≤ 0.01; *** = p ≤ 0.001.

Six proteins were found to be significantly increased in the conditioned media in the presence of IAV (**Fig. 4B**), all of which are involved in either evasion of host responses or modulation of bacterial virulence. Enolase and GAPDH are “moonlighting proteins,” serving more than one function in *Spn*[70]. In the cytosol they both function as glycolytic enzymes, however when present at the *Spn* surface, even in low abundance[71], they bind and activate host plasminogen increasing transmigration by degradation of the extracellular matrix[72–74]. In addition, enolase and GAPDH both inhibit complement activation by binding to either C4b-binding protein (C4BP)[75] or complement factor C1q[76], respectively. GAPDH binds to the complement factor C1q[76]. Pneumococcal surface adhesin A (PsaA) is a lipoprotein and a component of the membrane complex that transports manganese and zinc ions[77], and it is critical for *Spn* virulence[78, 79]. FusA, an elongation factor that is surface-associated in *Streptococcus oralis*[80], was significantly increased in the presence of IAV (**Fig. 4B**). Whether FusA is also a surface protein in *Spn* and functions in complement interference[81] needs further investigation. Overall, we found that IAV triggers the release of multiple *Spn* surface-associated proteins that mediate host invasion and subversion of host immune response (**Fig. 4C**). To further support the latter observation, we tested the ability of *Spn*’s condition media (after exposure to IAV) to modulate complement activity. Treatment of WT *Spn* with conditioned media from *Spn* challenged with IAV, we observed a decrease in phagocytosed bacteria by alveolar macrophages (MH-S) (**Fig. 4D**). We then tested for changes in complement hemolytic activity induced by *Spn* released proteins. Human serum was treated with concentrated conditioned media from an *Spn* deficient in pneumolysin (i.e. Δ*ply,* with/without exposure to IAV, to avoid inducition of cell lysis by the pore- forming activity of pneumolysin[82]), then serum was used to challenge sheep red blood cells. We observed that conditioned media from Δ*ply* exposed to IAV reduced the hemolytic activity of complement (**Fig. 4D**). Together, these results show that IAV can directly alter *Spn*’s ability to counter the effects of innate immune effectors.

### IAV leads to changes in pneumococcus cytotoxicity, capsule shedding, and bacterial adhesion though metabolic alterations

Our proteomic data demonstrated that IAV challenge led to decreased Ply expression in *Spn*. To further test how IAV alters the virulence of *Spn*, we first assessed how their interaction modulates *Spn-*induced toxicity in A549 type 2 respiratory epithelial cells. We observed that *Spn* challenged with IAV induced less cytotoxicity than an equivalent dose of *Spn* alone, comparable to that of a pneumolysin deficient mutant, Δ*ply* (**Fig. 5A**). These data further confirmed our previous results indicating reduced capsule production in *Spn* challenged with IAV (**Fig. 1B, Fig. S8**), we observed that IAV challenge induced capsule shedding in a dose dependent manner (**Fig. 5B**) and increased adhesion to A549 cells (**Fig. 5C**), in line with previous studies showing capsule shedding promotes host cell adhesion[83]. Increased adhesion was also observed in strains *Spn* D39, WU2 and 6A10 after bacterial incubation with IAV (**Fig. S11**). Interestingly, a significant reduction in cellular invasion was observed after *Spn* challenge with IAV (**Fig. 5D**). These results suggest that upon initial contact between *Spn* and IAV, the pneumococcus is primed towards an adhesion phenotype with low toxicity.

**Figure 5:**
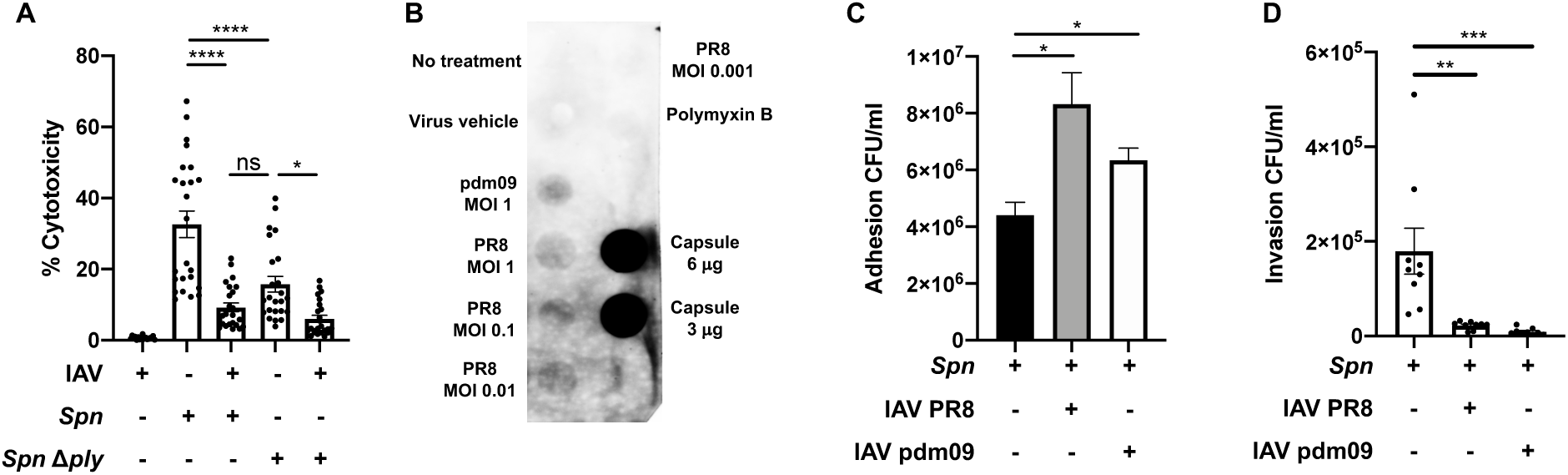
Influenza influences *Spn-*induced cytotoxicity and adhesion and promotes capsule shedding. (**A**) *Spn* alone is significantly more toxic to A549 cells than *Spn* challenged with IAV, or mutant *Spn* !1*ply*, as measured by LDH cytotoxicity assay. IAV continues to synergize with !1*ply* to further reduce toxicity. Data reflects n=24 shown as mean ± SEM analyzed by one- way ANOVA (*p<0.05, ****p<0.0001). (**B**) Dot blot of concentrated conditioned media from *Spn* with or without IAV challenge using two viral strains, PR8 and H1N1 A/California/7/2009 (pdm09). IAV challenge induced capsule release into media in a dose-dependent manner. Both IAV strains induced *Spn* to increase adhesion (**C**) and decrease invasion (**D**) of A549 cells. Data in (**C**) and (**D**) represent n=9 presented as mean ± SEM analyzed by Kruskal-Wallis test with Dunn’s multiple-comparison post-test. Asterisks denote the level of significance observed: * = p ≤ 0.05; ** = p ≤ 0.01; *** = p ≤ 0.001.

### IAV-driven *Spn* glucose and galactose metabolism modulates growth and virulence strategies

After translocation from the nasopharynx to the lung, availability of sugars such as glucose and galactose is significantly increased[84–87]. Our multiproteomic studies showed that IAV induced changes to glucose and galactose metabolism (**Fig. 2, 3, Fig. S9, 10**). To test the effect of glucose and galactose availability in the biological changes induced by IAV in *Spn*, we supplemented 50% Todd Hewitt Broth with either glucose or galactose at 1% w/v. IAV-induced reduction in pneumococcal growth was rescued by galactose but not glucose (**Fig. 6A**). We observed an increase in pneumolysin expression (**Fig. 6B**), toxicity (**Fig. 6C**) and activation of apoptosis (**Fig. 6D**) after glucose but not with galactose supplementation. While no changes in adhesion were induced by either sugar (**Fig. 6E**), significantly increased invasion was observed upon galactose supplementation (**Fig. 6F**). These results suggest that IAV-induced changes to *Spn* upon initial contact, are further modulated by metabolite availability.

**Figure 6:**
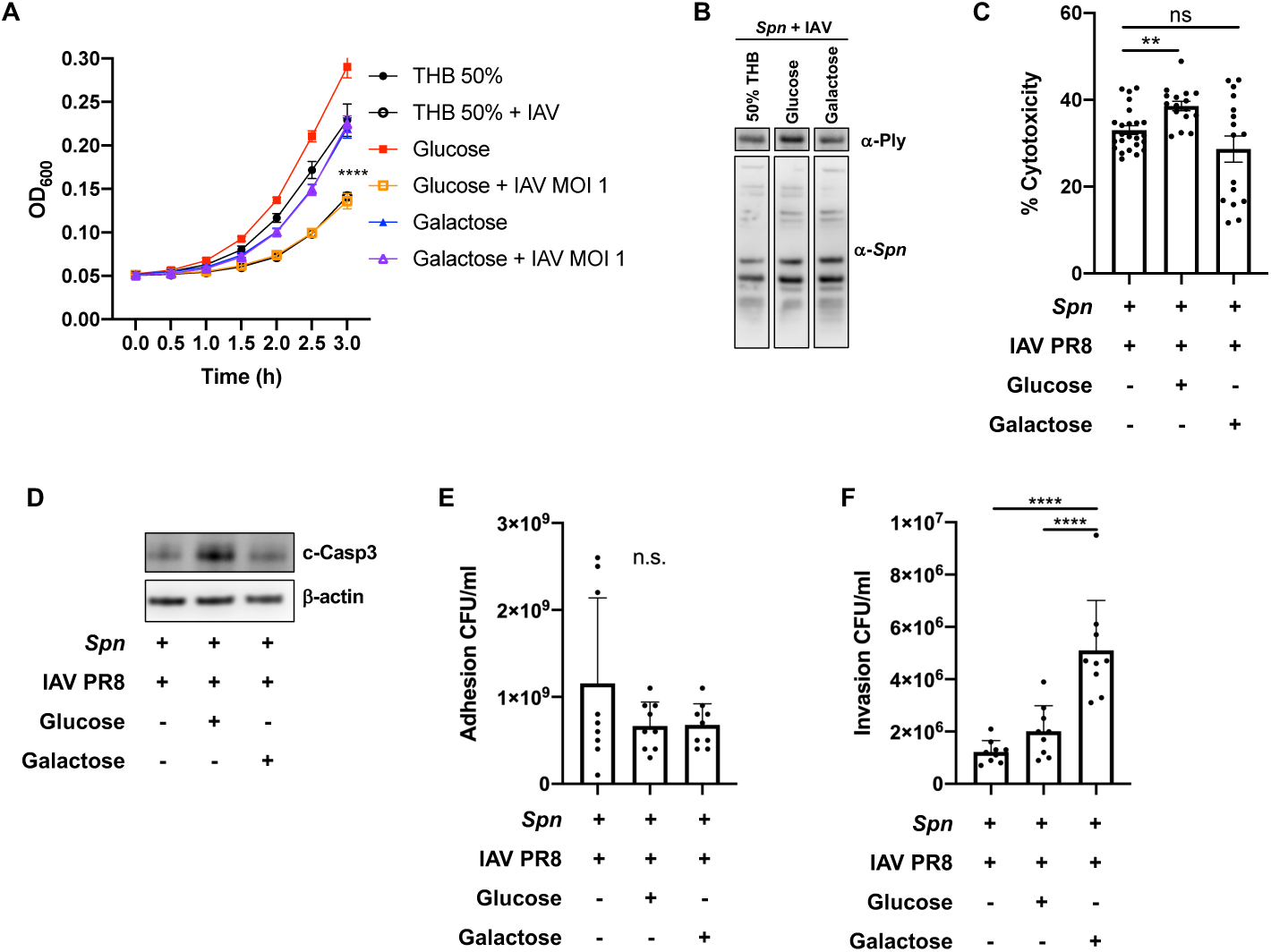
IAV-driven *S. pneumoniae* glucose and galactose metabolism modulates growth and virulence strategies. (**A**) Growth curves for Spn grown alone in minimal media (THB 50%, black closed circles), or supplemented with glucose 1% w/v (red solid squares) or galactose 1% w/v (blue solid triangles) do not differ significantly. Growth of *Spn* challenged with IAV was blunted when grown in minimal media (black open circles) or with glucose (orange open squares), but not with galactose (purple open triangles) Data reflect n=8 per condition (two-way ANOVA, ****p<0.0001 at t=3h). Glucose supplementation in media rescues (**B**) expression of pneumolysin as measured by Western blot, (**C**) exacerbated cytotoxicity to A549 cells as measured by LDH assay (n=8 per group, mean ± SEM, one-way ANOVA ****p<0.0001), and (**D**) cleaved caspase- 3 expression by A549 cells infected with *Spn*+IAV. (**E**) Adhesion of bacteria to A549 cells was unchanged regardless of sugar supplementation, but *Spn*+IAV supplemented with galactose (**F**) improved invasion into A549 over *Spn* grown in glucose or un-supplemented media. Statistical analyses in **E** and **F** reflect n=9 per condition, mean ± SD, analyzed by Kruskal-Wallis test with Dunn’s multiple-comparison post-test. Asterisks denote the level of significance observed: * = p ≤ 0.05; ** = p ≤ 0.01; *** = p ≤ 0.001.

### Influenza virus infection of host respiratory epithelial cells leads to additional alterations to the pneumococcal proteome

Colonizing *Spn* has been shown to translocate from the nasopharynx to the lung and cause severe disease after establishment of IAV infection. To evaluate changes in the proteomic profile of *Spn* after exposure to IAV infected type II respiratory epithelial cells (RECs), we used an *in vitro* model of infection, which was previously shown to predispose RECs to bacterial toxin mediated necroptosis [15]. Herein, we infected A549 respiratory epithelial cells with IAV PR8 at an MOI of 2 [15]. Four hours post infection, cells were challenged with *Spn* at an MOI of 10 for an additional four hours. As controls *Spn* was grown in tissue culture media with or without IAV exposure for the duration of the experiment. A total of 369 proteins were expressed differentially (P < 0.05) (**Fig. 7A**). Principal Component Analysis (PCA) showed differential clustering of the bacterial proteins, suggesting each growth condition had an effect in pneumococcal biology (**Fig. 7B**). Hierarchical clustering revealed 3 distinct clusters of *Spn* proteins that changed expression between groups (**Spreadsheet S6**). Evaluation of the biological processes (GO terms) associated with cluster 1 differentially expressed proteins revealed mainly a decrease in proteins associated with cell wall biogenesis, nucleoside triphosphate biosynthetic process, peptidoglycan metabolic process, regulation of cell morphogenesis, and ion transport (cluster 1, **Fig. 7C**). In cluster 2, proteins showed increased expression only in the *Spn* isolated from the A549 cells that were previously infected with IAV. For these proteins GO terms associated with cell septum assembly, cytokinetic process, ATP biosynthetic process, and purine nucleoside triphosphate biosynthetic process (cluster 2, **Fig. 7C**). Lastly, proteins in cluster 3 revealed mainly an increase in proteins involved in 5- phosphoribose 1-diphosphate biosynthetic process, organophosphate metabolic process and tRNA aminoacylation, relative to unchallenged *Spn* control (cluster 3, **Fig. 7C**). Interestingly, *Spn* proteins upregulated in the presence of the influenza infected host showed a shift in purine metabolism, found decreased upon direct interaction between *Spn* and IAV.

**Figure 7:**
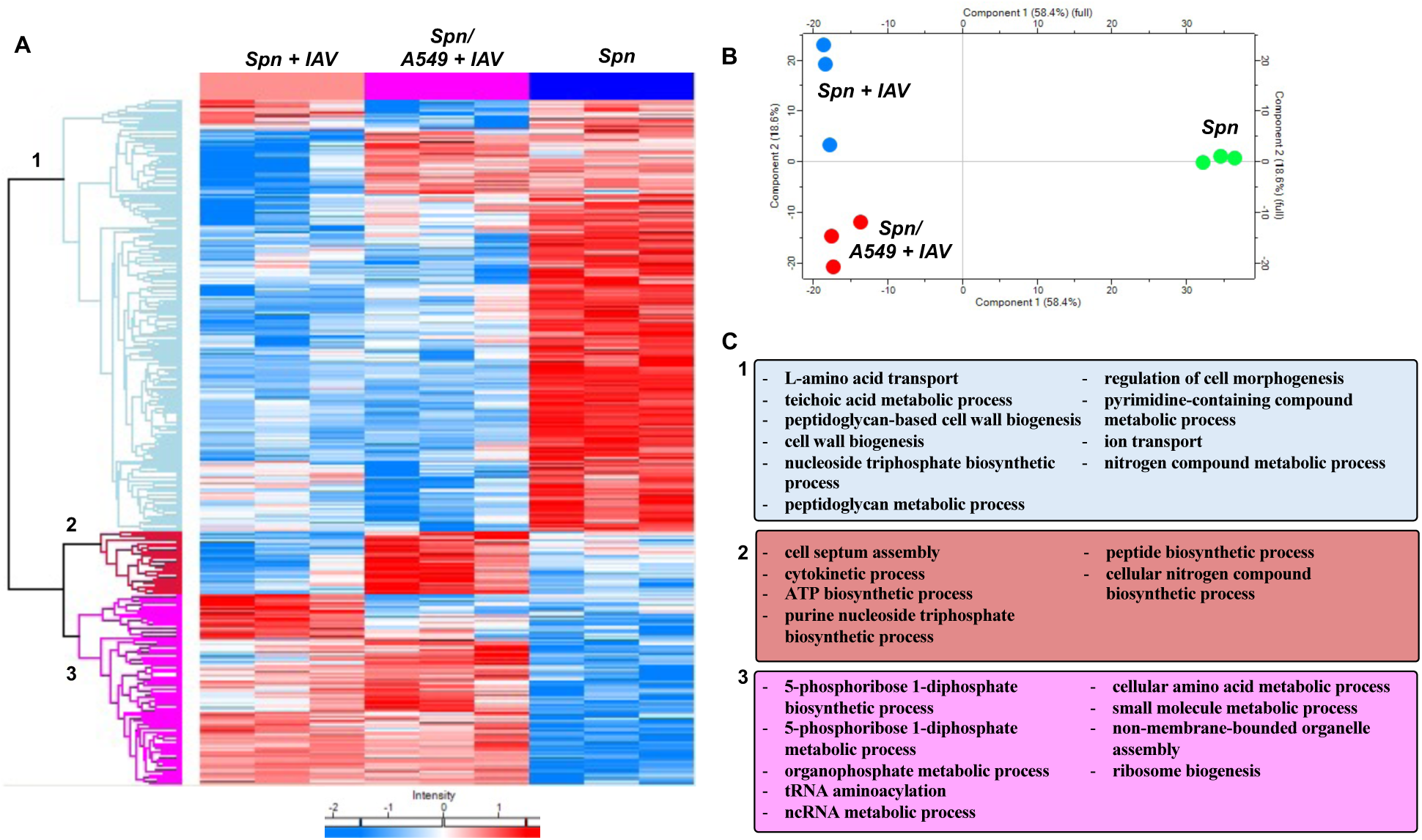
Primary IAV infection of respiratory epithelial cells leads to additional alterations to the pneumococcal proteome. A549 respiratory epithelial cells were infected with IAV PR8 at an MOI of 2. Four hours post infection, cells were challenged with *Spn* at an MOI of 10 for an additional four hours. As controls *Spn* was grown in tissue culture media with or without IAV exposure for the duration of the experiment. (**A**) Proteomic changes of *Spn* after IAV challenge, in the presence of IAV infected A549 cells or mock challenge. Hierarchical clustering of label-free quantification (LFQ) intensities of significantly changed proteins (ANOVA, P,0.05) revealed 3 distinct clusters. Their abundance profiles among the groups were plotted in the heatmap (**A**). (**B**) Principal component analysis denoting differences in *Spn* proteomic profiles. (**C**) Enriched GO biological process terms are indicated for each marked cluster.

### Sequential infection of RECs with IAV and *S. pneumoniae* leads to differential proteomic remodeling

Here we evaluated changes in the proteomic profile of RECs after sequential infection with IAV and *Spn* as previously described [15]. A549 cells were mock challenged or infected with IAV, *Spn* or sequentially infected with these pathogens. A total of 315 proteins were expressed differentially (P < 0.05) between the groups (**Fig. 8A**). Principal Component Analysis (PCA) showed that infected cells had significant proteome remodeling when compared to mock (vehicle) infected cells (**Fig. 8B**). Hierarchical clustering revealed 3 distinct clusters of A549 proteins that changed expression between groups (**Spreadsheet S7**). Evaluation of the biological processes GO terms associated with cluster 1 differentially expressed proteins revealed in its majority an increase in proteins associated with RNA export from nucleus, ribonucleoprotein complex localization, peptidoglycan metabolic process, MRNA splicing, via spliceosome, and Intracellular transport (cluster 1, **Fig. 8C**). In cluster 2, proteins showed both increased and decreased expression, GO terms associated with these proteins were ATP metabolism, generation of precursor metabolites, energy derivation by oxidation of organic compounds, and viral processes (cluster 2, **Fig. 8C**). Lastly, proteins in cluster 3 mainly increased in expression and were found to be involved in receptor binding, lipoprotein clearance, and receptor-mediated endocytosis involved in cholesterol transport (cluster 3, **Fig. 8C**). These results suggest that viral and bacterial infections drive global changes to REC proteome, however upon sequential infection, the response is similar to the single bacterial infection group.

**Figure 8:**
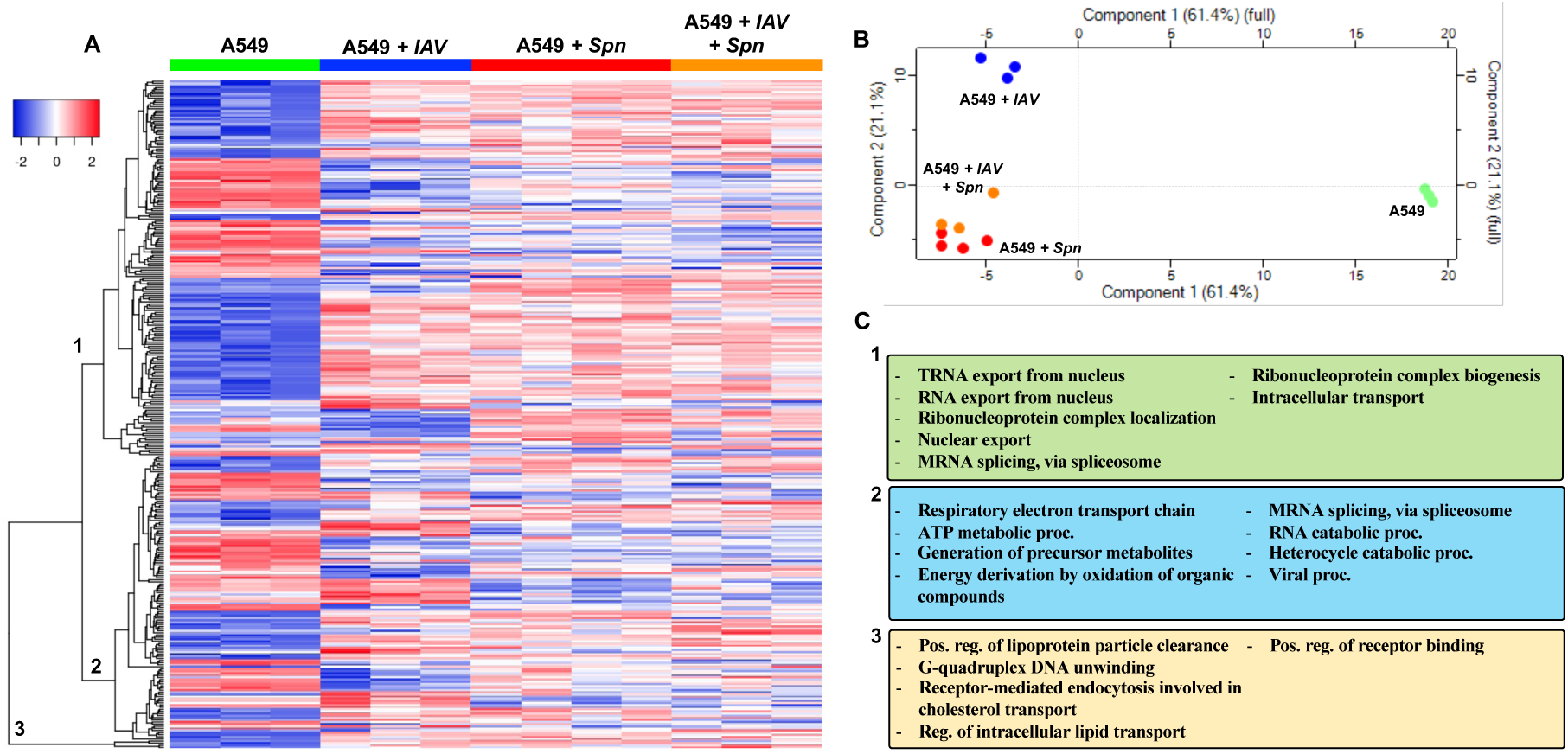
Sequential infection of RECs with IAV and *Spn* leads to differential proteomic remodeling. A549 respiratory epithelial cells were infected with IAV PR8 at an MOI of 2. Four hours post infection, cells were challenged with *Spn* at an MOI of 10 for an additional four hours. As control A549 was grown in tissue culture media exposure to any pathogen. Proteomic changes in A549 cells. (**A**) Hierarchical clustering of label-free quantification (LFQ) intensities of significantly changed proteins (ANOVA, P, 0.05) revealed 3 distinct clusters. Their abundance profiles among the groups were plotted in the heatmap. (**B**) Principal component analysis denoting differences in *Spn* proteomic profiles. (**C**) Enriched GO biological process terms are indicated for each marked cluster.

## Discussion

Recent research has demonstrated direct interaction between nasopharyngeal commensal bacteria and IAV, causing a synergistic increase in susceptibility to bacterial infection and disease severity[88]. In this study, we define the biological alterations IAV induces upon direct interaction with *Spn.* We used the power of multi-proteomic approaches to reveal candidates for the underlying molecular interactions between *Spn* and IAV. Using AE-MS, we found *Spn* proteins that interact with IAV. Future experiments using other *Spn* mutants lacking putative IAV- interacting genes should be performed to validate candidate interaction partners in detail. Our experiments using *Spn* lacking the candidate IAV-interacting protein pneumolysin (Δ*ply*) demonstrated that IAV affects multiple molecular pathways in *Spn* (regarding host cytotoxicity), confirming data from our proteomic analyses. The combined global and phosphoproteome analyses of *Spn* challenged with IAV revealed themes of altered metabolism, including glycolysis, purine/pyrimidine synthesis, amino acid synthesis and a switch to galactose metabolism. Galactose metabolism is an alternative energy production pathway that has been linked to *Spn* virulence[44, 45]. Proteins that mediate cell division are more abundant (Fts proteins) or increased in phosphorylation (StkP and substrates), suggesting a possible effort by the bacteria to modulate cell growth without undergoing autolysis. Importantly, the slow growth of *Spn* early after contact with IAV may be a way for bacteria released from the nasopharynx to prevent initial immune activation and evade the effect of host antimicrobials while colonizing the respiratory tract[89–91]. It remains to be determined if, upon *in vivo* infection, these metabolic changes will lead to faster growth due to the abundance of galactose in the respiratory tract[92]. This suggests IAV directly primes *Spn* towards metabolic fitness to promote pulmonary infection.

The observed changes in the proteome and phosphoproteome suggested a possible switch of gene expression towards an infectious phenotype. Also, the *Spn* biofilm dispersal observed upon contact with IAV also support virulence switch as planktonic but not biofilm are associated with pulmonary and invasive disease[93]. While the adhesion and immune evasion factor PspA was up-regulated in the presence of IAV, IgA and Ply were down-regulated. It was previously shown that mice challenged with pneumolysin have reduced influenza-associated pathogenesis[94], which leads us to speculate that to counteract this effect, IAV may down- regulate the production of Ply. PspA upregulation by IAV may also promote its binding to host lactoferrin, potentiating iron acquisition and bacterial growth in the lower respiratory tract[95]. It also indicates that IAV-mediated protein regulation may be under a more sophisticated spatial and temporal control that fine tunes *Spn* virulence depending on the infection stage.

Capsule is known as an important virulence factor of *Spn* and is the basis for commercial vaccines. While capsule is required for bacterial survival in the bloodstream (invasive pneumococcal disease) and immune evasion[96], its reduction is not completely detrimental for the bacterium’s ability to cause respiratory disease[97] and cardiac damage[98]. Non-typeable *Spn* lacking capsule also exists and has been shown to be equally pathogenic[99]. Moreover, capsule shedding was recently described as a new pathway for the development of invasive disease[83]. We investigated whether IAV challenge alters the association of *Spn* surface proteins due to the observed reduction in capsule. Indeed, several *Spn* proteins, some previously known to be surface attached, were found to be increased extracellularly in the presence of IAV. It is unlikely that these proteins were released by active secretion, since *Spn* only has a Type II system that is used to transform DNA[100, 101], and there is no evidence suggesting that the Sec translocase pathway is involved in surface localization of the identified *Spn* proteins[102, 103]. Whether additional mechanisms exist to facilitate the release of *Spn* surface proteins, or if they are simply untethered from *Spn* surface due to capsule shedding, needs to be further studied. However, the significantly increased release of enolase, GAPDH, and PsaA that can facilitate plasminogen and complement binding and modulate host immune response suggests that capsule shedding is another strategy *Spn* uses to promote virulence upon contact with IAV. Of interest, while the majority of the proteomic changes caused by direct influenza virus were observed when infecting a system were epithelial cells were present, a set of bacterial proteins were further changed suggesting further adaptation as response to host signaling.

How pathogens adapt and tune their virulence in the polymicrobial environment is a fundamental question that needs to be addressed to truly understand pathogen behavior in the host. Pulmonary infections are commonly caused by respiratory pathogens including viruses and bacteria, and in many circumstances, a combination of both. Our results provide insights into the molecular mechanism behind the early stages of *Spn*-IAV pathogenic synergy. This may directly translate to bacterial synergism with SARS-CoV-2 viral infection[104, 105], providing another possible mechanism for development of severe infections. We foresee our results being directly applicable to studies in other bacteria-virus interactions including other nasopharyngeal commensals and pathogenic viruses.

## Materials and Methods

### Bacterial and viral cultures

*Spn* serotype 4 strain TIGR4 was grown overnight on 1% agar- Todd Hewitt Broth (Thermo Fisher Scientific) with yeast (THY) plates supplemented with catalase. Liquid cultures of THY were inoculated and grown to log-phase, then influenza A virus strain Influenza A/Puerto Rico/8/1934 (PR8) was added (1:1 ratio) and incubated for 1 h. Selected experiments used influenza A virus strain under the same conditions. *Spn* grown under the same condition without added IAV served as control, and each experiment was performed with three biological replicates. PR8 was propagated in embryonated chicken eggs[106, 107]. Stock viruses were titered using plaque assays on MDCKs. IAV was heat-inactivated at 56°C for 30 min[108]. Growth curve was done by first growing *Spn* to log phase in THY, then seeded in a 96-well plate at a volume of 100uL with/without IAV or heat-killed virus (HK IAV) and OD600 was measured every 30 minutes. Transparent/opaque phenotype assessment was done as previously described[98]. For experiments regarding the effects of glucose and galactose in the IAV-induced bacterial changes, bacteria were grown in 50% THB with either 1% w/v glucose or galactose or without sugar supplementation[109]. Bacterial cells were quantified upon growth in each medium and diluted to an MOI of 10 for infection of A549 respiratory epithelial cells (ATCC). Adhesion and invasion assays were done as previously described[98, 110]. For transcriptomics analysis TIGR4 was grown in CY media to around OD600=0.4. 100ul of PR8 stock at 10^8^ TCID50/ml was added to 10ml of TIGR4 cultures. For negative controls, 100ul of mock was added to 10ml of TIGR4 cultures. Cultures were incubated turning end-over-end at 37°C for 15, 30, or 60 minutes. Each time point and condition (virus/mock) was performed in triplicate. After incubation, cultures were immediately spun down at 6,000xg and supernatant was discarded. Pellets were resuspended in 500ul RNAprotect Bacteria Reagent (QIAGEN, mat. 1018380) and frozen at -80°C. Isogenic mutant deficient in ply were created by insertion of an erythromycin resistance cassette, ermB, using allelic exchange, and grown in medium supplemented with 1 μg/ml of erythromycin as previously described[111, 112]. Co-incubation model also include the use of *E. coli, P. aeruginosa* and *S. marcescens*[22, 23] following the same protocol described above for *Spn*.

### Cell culture

A549 type II alveolar epithelial cells were infected with IAV at MOI 2 for 4 hours, and subsequently challenged with *Spn* at an MOI 10 for 4 hours as recently described[15].

### Immunoblots

Samples for Western blotting were homogenized in PBS and sonicated, then diluted in Laemmli buffer and aliquoted for storage at -80 degrees C. Samples were loaded into 4-15% gradient gels at 20 μg per lane, separated by SDS-PAGE, and transferred to nitrocellulose membranes. Total protein was quantified by Ponceau stain, then membranes were blocked in blocking buffer (TBS-0.01% Tween-20 containing 5% BSA) for at least 1 h. Primary antibodies were diluted in blocking buffer and membranes were incubated overnight at 4 degrees C. Primary antibodies used included: anti-pneumolysin antibody (ab71810, Abcam), anti-*Spn* serotype 4 (#16747, Statens Serum Institut) and cleaved-caspase-3 (AF835SP, R&D Systems) and β-actin as loading control (ab8226, Abcam). HRP-tagged secondary antibodies were used to detect primary antibodies (1:10,000 in blocking buffer) for 1 h. SuperSignal West PICO Plus (ThermoFisher 34580) was used to develop HRP. All images were collected on an Amersham Imager 680 (GE) and analyzed for densitometry in ImageJ. Briefly, to relatively quantify protein bands from western blots relative to loading control (herein, total loaded protein).

### Capsule Shedding Assay

TIGR4 bacteria was grown to OD600 0.5 in THY, washed once with SMM buffer (0.5M sucrose, 0.02M MgCl2, 0.02M MES, pH 6.5), and resuspended in 1/10 volume SMM buffer. Bacteria were incubated with IAV strains (MOI 1-0.001) or polymyxin B (32 mg/ml) for 30 minutes at 37°C, then spun out at high speed. Supernatant was treated with proteinase K for 30 minutes at 37°C, then spotted on nitrocellulose membrane alongside serotype 4 capsular polysaccharide 3-6 μg (SSI Diagnostica cat. 76855). The membrane was then blocked with 5% BSA in TBS with 0.05% Tween-20 and probed overnight with anti-pneumococcus capsular antibody (1:500, SSI Diagnostica cat. 16747). Membranes were washed extensively, then probed with HRP-conjugated anti-rabbit secondary antibody and developed with Pierce ECL Western blotting substrate (ThermoFisher cat. 32109).

### Cell lysate preparation, and protein digestion

The fresh cell pellets were first rinsed with cold PBS to remove culture medium contamination. The pellet fraction was then lysed with 2 x SED lysis buffer (4% SDS, 50mM EDTA, 20 mM DTT, 2% Tween 20, 100mM Tris-HCl, pH 8.0) followed by sonication (Misonix 3000, Ultrasonic Cell Disruptor) at amplitude 6 in six 30 s on/off cycles while cooling the lysates in an ice-water bath. After a 10 min centrifugation step at 16,000 × g, the soluble lysate fraction was collected. To estimate the protein concentration, an aliquot (10 ul) of each sample was analyzed on an SDS-PAGE gel alongside a known amount of BSA (2 μg)[113]. Proteins were digested following a Suspension Trapping (STrap) protocol with the self-packed glass fiber filters (Whatman, GF/F)[30]. After digestion, the resulting peptides were dried in a SpeedVac (Thermo Scientific) followed by C18-based desalting using StageTip protocol[114]. The peptides were lyophilized and then stored in -80°C until further analysis.

### AE-MS procedure

Log-phase grown *Spn* (∼1 × 10^11^ cells) were harvested, washed with PBS, and lysed by sonication in PBS containing HALT protease inhibitor cocktail (#78430, ThermoFisher). Clear lysate with soluble *Spn* proteins was obtained by centrifugation at high speed for 15 min, and half of it was mixed with 1 × 10^8^ IAV and 40 μL anti-HA agarose beads (Pierce cat. #26181) while the other half was directly mixed with anti-HA beads (control). Each group was performed in 3 replicate experiments. The mixing was performed overnight at 4°C with rotation. The next day, the beads were settled by centrifugation, the lysate was removed, and the beads were washed extensively with PBS containing 0.05% Tween 20. Viral particles and *Spn* proteins enriched on the beads were eluted by 5% SDS in 50 mM triethylammonium bicarbonate (TEAB) and digested using the STrap method[30, 115]. After digestion, peptides were desalted using the established StageTip method as descried above, and stored in -80⁰C until further analysis.

### *Spn* phosphopeptide enrichment

Around 200 μg peptides resulted from STrap digestion of *Spn* culture with or without IAV co-incubation were subjected to enrichment using titanium dioxide beads according to the established protocol[57]. After enrichment, peptides were desalted using StageTips before LC-MS/MS analysis.

### *Spn* conditioned media collection

*Spn* was grown as described above to log-phase, then incubated with IAV (or vehicle control) at a 1:1 ratio for 1 h at 37°C in quadruplicate. Cultures were harvested by centrifugation at 9,000 × g for 15 min, and the supernatant was collected and filtered through a 0.45 μm disc filter. Proteins in the supernatant were concentrated by Amicon Ultra-15 filter (10K MWCO) and buffer exchanged into 50 mM ammonium bicarbonate by centrifugation. Concentrated proteins were collected and mixed with 4% SDS and 25 mM DTT, then digested using the STrap method as described above. Conditioned media was also used for macrophage phagocytosis assay (using MH-S cells)[21, 116] and complement hemolysis assays (using sheep red blood cells) [117].

### LC-MS/MS analysis

The LC-MS/MS analysis was conducted on an Ultimate 3000 RSLCnano System coupled to a quadrupole Orbitrap mass spectrometer (Q Exactive, Thermo Scientific) via a nano electrospray ion source. The experimental methods were described previously in detail[118] In brief, the desalted peptide samples were first dissolved into 20 μl LC buffer A (0.1% formic acid in water), and then loaded onto a PepMap C18 trap column (100 μm x 2 cm, 5 μm particle size, Thermo Scientific) followed by separation on an in-house packed column (75 mm x 19 cm, 3.0 mm ReproSil-Pur C18-AQ). Depending on sample complexity, we utilized varied LC gradient to maximize the identification rate and throughput. For the host cell (A549) related experiment, a 220-min gradient (180 min from 2% to 35% buffer B, 0.1% formic acid in acetonitrile; 10 min to 80% B) was used. For the samples from pulldown assay, phosphorylation enrichment, and Spn/Flu experiment, a 150-min gradient (120 min from 2% to 35% buffer B, 0.1% formic acid in acetonitrile; 10 min to 80% B) was used. The full (MS1) scans were acquired at a resolution of 70,000 with a mass range of *m/z* 300-1,700 in a data-dependent mode. The ten most intense ions were selected from each cycle for MS/MS (MS2) analysis using high-energy collisional dissociation (HCD) with normalized collision energy of 27%. The dynamic exclusion time was set to 20 seconds. Singly charged ions and ions with five or more charges were excluded for MS/MS analysis. The MS2 scans were performed at a resolution of 17,500. Target value for the full scan MS scan was 5 × 10^5^ with a maximum injection time of 20 ms, and for MS/MS scan was 1 × 10^6^ with a maximum injection time of 100 ms.

### Protein identification and quantitation

The MS raw data were searched against a meta database that contained protein sequences of *Streptococcus pneumoniae* serotype 4 (strain ATCC BAA-334 / TIGR4) (UniProt taxon identifier 170187; 2,115 sequences), Influenza A virus (strain A/Puerto Rico/8/1934 H1N1) (UniProt taxon identifier 211044; 112 sequences) and Homo sapiens (UniProt taxon identifier 9606; 17,023 reviewed sequences). To determine proteins associated with culture media additional searches for Canis Lupus familiaris (UniProt taxon identifier 9615), Bos taurus (UniProt taxon identifier 9913) and Sus scrofa domesticus (UniProt taxon identifier 9825) were performed. The MaxQuant-Andromeda software suite (version 1.6.5.0) was employed with most of the default settings[119]. Two mis-cleavage sites were allowed; the minimum peptide length was set to seven amino acids. Protein N-terminal acetylation and oxidation (M) and were set as variable modifications; carbamidomethylation (C) was set as fixed modification. The MS1 and MS2 ion tolerances were set at 20 ppm and 10 ppm, respectively.

The false-discovery rate (FDR) of 1% was set at both peptide and protein level. The embedded label-free algorithm (MaxLFQ) was enabled using a minimum ratio count of 1. Further bioinformatics analyses such as Hierarchical clustering, Principal Component Analysis (PCA), t- tests, volcano plots and correlation analyses were performed in Perseus software (version 1.6.7.0)[120]. The LFQ data were first log2 transformed and then filtered to include those that were present in at least two of the three biological replicates in one of the two groups. The missing values were imputed based on default parameters. For hierarchical clustering, the Z-scored LFQ intensities were used with Euclidean as a distance measure for both column and row clustering.

### RNA Extraction

TIGR4 samples were thawed and pelleted at 6,000xg for 10 minutes. Supernatant was discarded. Pellets were resuspended in 400μl of RLT Lysis Buffer (QIAGEN mat. 1015762). 10μl of 2-Mercaptoethanol had been added to the RLT buffer for every 1 ml buffer. All 400μl samples were loaded into Lysing Matrix E tubes (MP Biomedicals, ref. 6914100). Samples were lysed with the Fast Prep-24 (MP Biomedicals) homogenizer for 10 pulses, 20 seconds each. All samples were incubated at 70°C for 10 minutes or until clear. Lysate was spun through QIA Shredder columns (QIAGEN, cat. 79656) for 1 min at max speed. Flow-through was returned to the column and spun again for a total of three spins. Samples were combined with 400μl of 70% ethanol and loaded onto RNeasy mini columns (QIAGEN RNeasy Mini Kit, cat. 74104). RNA was extracted following the RNeasy protocol, eluting in 70μl of nuclease-free water. RNA sample concentrations were measured by Nanodrop. RNA-quality was assessed by the Hartwell Center for Biotechnology at St. Jude Research Hospital.

### DNA Removal

For each TIGR4 RNA sample, 10μg RNA was combined with 1μl rDNAse I from the DNA Removal Kit (Thermo Scientific cat. AM1906). A tenth of the total volume of 10X DNase I buffer was added to each sample. Samples were incubated at 37°C for 30 minutes. 2μl DNase Inactivation Reagent was added to each sample. Samples were incubated at RT for 2 minutes. Samples were spun at 10,000xg for 90 seconds. Supernatant was transferred and measured by Nanodrop.

### Ribosomal RNA Depletion

The ribosomal RNA depletion protocol was adapted from Culviner et al. 2020[121]. Biotinylated primers were combined into a mix to pull down rRNA. The mix contained 10μl of each 5S primer and 5μl of the remaining primers. Primers started at the stock concentration of 100μM. 2X Binding/Wash (B&W) buffer was made to contain 10 mM 7.6 pH Tris, 1mM EDTA, and 2M NaCl. For each 10 samples, Dynabeads™ MyOne™ Streptavidin C1 beads (Thermo Scientific, cat. 65002) were prepared by pulling down 1.41ml beads with a magnetic rack. Beads were washed 3 times with 1.5ml 1X B&W buffer before being resuspended in 300μl 2X B&W. 10μl SUPERase-IN™ RNase Inhibitor (Thermo Scientific, cat. AM2696) was added to each batch. 30μl of prepared beads were used for each sample. Each sample mix received 3μl 20X SSC, 1μl 30mM EDTA, 1μl diluted oligo mix, 3μg of RNA sample, and water up to 30μl in a PCR tube. Samples were incubated at 70°C for 5 min in a thermocycler, stepping down a degree every 30 seconds down to 25°C. Samples were combined with the 30μl of beads and were thoroughly mixed. Tubes were incubated for 5 minutes RT, vortexed, and incubated again for 5 minutes at 50°C. Beads were pulled down by magnet and supernatant was transferred to new tubes. Each sample received 140μl water, 20μl 3M NaOAc pH 5.5, 2μl GlycoBlue™ (Thermo Scientific, cat. AM9516), and 600μl pure ethanol. Samples were kept at -20°C. Samples were spun at 14,000xg at 4°C for 30 minutes. Supernatant was removed; pellets were washed with 800μl cold 70% ethanol. Samples were spun at 14,000xg for 5 minutes at 4°C. Supernatant was discarded; pellets were resuspended in 10μl nuclease-free water and stored at -20°C. Used primers are shown in **Table S2**.

### Generating cDNA

NEB Next Ultra II RNA Library Prep Kit for Illumina (#E7775) was used to generate cDNA from 5μl RNA sample according to the provided protocol using the random primer method. Products were purified with Agencourt AMPure XP beads (Beckman Coulter Product No. A63881). 144μl resuspended beads were added to 80μl product, incubating at RT for 5 minutes. Beads were pulled down by magnet, supernatant was removed. Beads were washed twice with 200μl 80% ethanol. Beads were left at RT to dry 5 minutes before eluting with 53μl TE. 50μl supernatant was transferred to new plate.

### Adaptor Ligation

NEB Next Ultra II RNA Library Prep Kit for Illumina was used to ligate adaptors to 50μl samples of cDNA. Products were purified by AMPure XP beads as previously described, eluting in 17μl 0.1X TE. 15μl supernatant was transferred to a new plate. PCR enrichment for adaptor ligated DNA was also performed using NEB Next Ultra II RNA Library Prep Kit for Illumina. AMPure XP beads were used to purify PCR products, eluting in 23 μl 0.1XTE, transferring 20μl to another plate for sequencing.

### Sequencing

The library was prepped using the TruSeq Stranded Total RNA by Illumina (PN. 20020597). Samples were sequenced on the Illumina Novaseq 6000.

### RNA-seq analysis

Raw paired-end reads in FASTQ format were processed by fastp v0.20.0 to remove adaptor sequences, short reads (length < 20) and low quality reads with a Phred quality score < 25. The resulting clean reads were aligned against the respective *Spn* reference genomes (TIGR4: NC_003028.3, D39: NC_008533.2, BHN97x: CP025076.1) using bowtie2 (version 2.3.5) based on different experimental designs. SAM/BAM files were processed by Samtools 0.1.19. The featureCounts v2.0.1 from the Subread package was used to count mapped reads for genes with a MAPQ value ≥ 10. Raw read counts were first normalized by variance stabilizing transformation by DESeq2, and then differential gene expression analysis was carried out by DESeq2 (log2-fold ≥1 and adjusted p-values ≤0.05). Heatmaps and volcano plots were generated to visualize statistically significant genes by pheatmap (1.0.12) and EnhancedVolcano (1.4.0), respectively, in R version 3.6.3.

### Data considerations and statistics

Unless otherwise noted, all *in vitro* experiments are composed of a minimum of 3 biological replicates, with ≥3 technical replicates each. Statistical comparisons were calculated using GraphPad Prism 8 (La Jolla, CA). Comparisons between two cohorts at a single time point are calculated by Mann-Whitney U test. Comparisons between groups of >2 cohorts or groups given multiple treatments were calculated by ANOVA with Tukey’s (one-way) or Sidak’s (two-way) post-test or by Kruskal-Wallis H test with Dunn’s multiple comparison post-test, as determined by the normality of data groups. Repeated measures are accounted for whenever applicable.

### Data Sharing

The mass spectrometry proteomics data have been deposited to the ProteomeXchange Consortium via the PRIDE partner repository with the dataset identifier PXD016214 and PXD016122. The raw RNA-seq data has been deposited to NCBI database under the accession number PRJNA837613.

## Declarations

### Ethics approval and consent to participate

Not applicable

### Consent for publication

Not applicable

### Availability of data and materials

The datasets generated and/or analyzed during the current study are available from the corresponding author on reasonable request.

### Author Contributions

M.P.P., Y-H.L., T.P., I.V., R.W-R, and Y.Y. carried out the experiments, M.P.P., Y-H.L., Y.Y. J.W.R. and N.G.J contributed to the design and conceptualization of the project. M.P.P., Y-H.L., C.A.M., J.W.R., Y.Y. and N.G.J wrote and edited the manuscript.

## Acknowledgements

Not applicable

