## Supplemental Figures for "A multiomics analysis of direct interkingdom dynamics between influenza A virus and *Streptococcus pneumoniae* uncovers host-independent changes to bacterial virulence fitness"

**Supporting Information:**

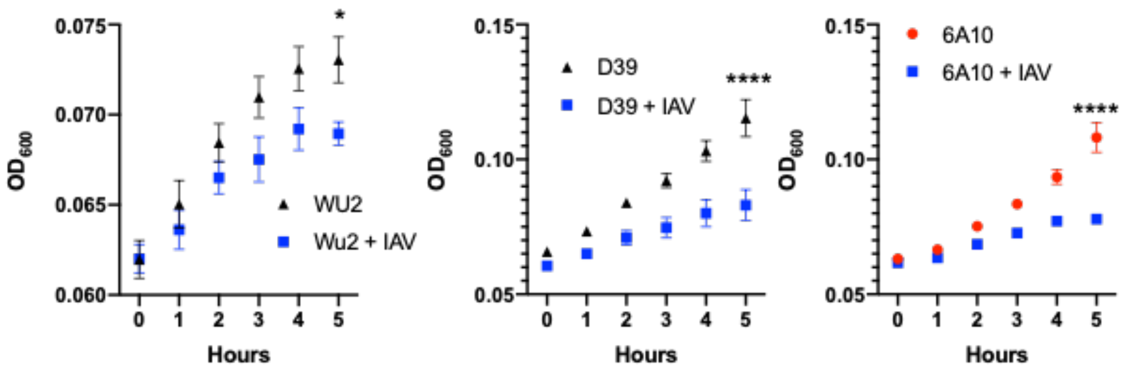


**Figure S1: IAV directly exerts effects on *Spn* independent of serotype.** Growth curve of *Spn* strains WU2, D39 and 6A10 in liquid media with/without IAV (H1N1 A/California/7/2009 [pdm09]). Data points represent mean +/- SD (two-way ANOVA, **p<0.05, ***p<0.0001 at t=5h).

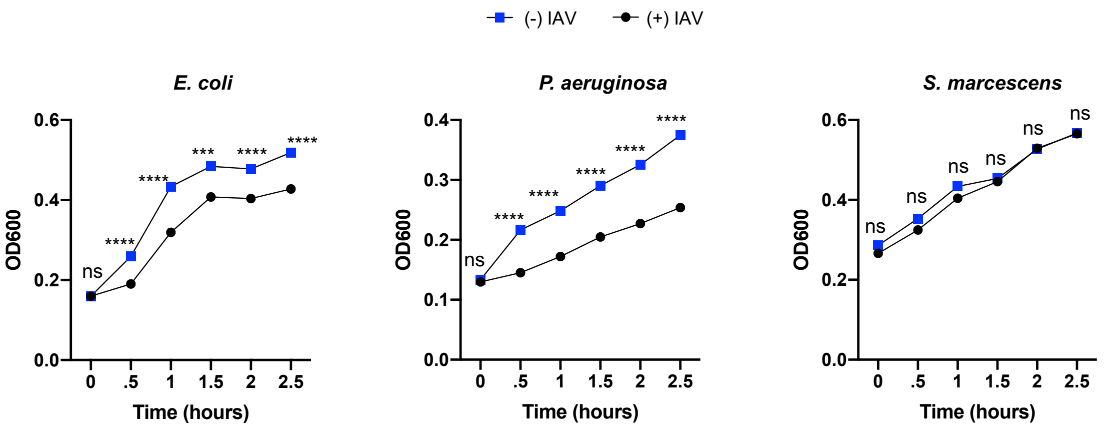


**Figure S2: IAV directly exerts effects on *E. coli* and *P. aeruginosa* but not *S. marcescens*.** Growth curve of *E. coli, P. aeruginosa* and *S. marcescens* in liquid media with/without IAV (PR8). Data points represent mean +/- SD (two-way ANOVA, *p<0.05, **p<0.01, ***p<0.001 and ****p<0.0001).

A

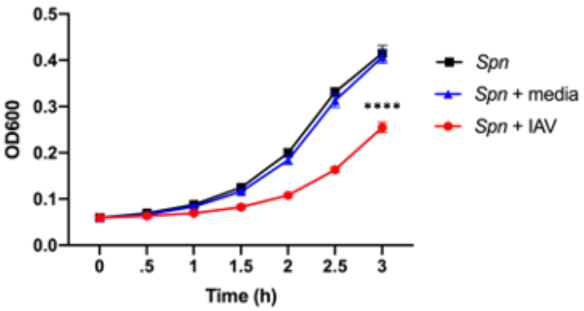


B

| Accession | Description | # PSMs | # Peptides | MW [kDa] |
|---|---|---|---|---|
| E2R0F2 | Filamin B OS=Canis lupus familiaris OX=9615 GN=FLNB PE=3 SV=2 | 866 | 47 | 281 |
| P02769 | SWISS-PROT:P02769 (Bos taurus) Bovine serum albumin precursor | 3052 | 39 | 69.2 |
| F1PWW0 | Filamin A OS=Canis lupus familiaris OX=9615 GN=FLNA PE=3 SV=3 | 684 | 38 | 280.5 |
| F1MRC2 | Myosin-2 OS=Bos taurus OX=9913 GN=MYH2 PE=3 SV=1 | 1060 | 37 | 223.2 |
| Q9BE40 | Myosin-1 OS=Bos taurus OX=9913 GN=MYH1 PE=2 SV=2 | 753 | 35 | 222.9 |
| O75369 | Filamin-B OS=Homo sapiens OX=9606 GN=FLNB PE=1 SV=2 | 655 | 35 | 278 |
| F1MM07 | Myosin-7 OS=Bos taurus OX=9913 GN=MYH7 PE=3 SV=3 | 678 | 33 | 223.1 |
| P49824 | Myosin-7 OS=Canis lupus familiaris OX=9615 GN=MYH7 PE=1 SV=3 | 686 | 32 | 222.8 |
| A0A0B8RSX6 | Filamin A, alpha OS=Sus scrofa domesticus OX=9825 GN=FLNA PE=3 SV=1 | 610 | 32 | 279.7 |
| P13533 | Myosin-6 OS=Homo sapiens OX=9606 GN=MYH6 PE=1 SV=5 | 657 | 31 | 223.6 |
| F1SS64 | Myosin-4 OS=Sus scrofa OX=9823 GN=MYH4 PE=3 SV=2 | 652 | 30 | 223.9 |
| P02453 | Collagen alpha-1(I) chain OS=Bos taurus OX=9913 GN=COL1A1 PE=1 SV=3 | 5246 | 27 | 138.9 |
| A0A5G2QQE9 | Collagen type I alpha 1 chain OS=Sus scrofa OX=9823 GN=COL1A1 PE=1 SV=1 | 2728 | 25 | 140.8 |
| P02452 | Collagen alpha-1(I) chain OS=Homo sapiens OX=9606 GN=COL1A1 PE=1 SV=5 | 2860 | 24 | 138.9 |
| P03466 | Nucleoprotein OS=Influenza A virus (strain A/Puerto Rico/8/1934 H1N1) OX=211044 GN=NP PE=1 SV=2 | 3039 | 22 | 56.2 |

**Figure S3. IAV directly exerts effects on *Spn* independent of viral stock source.** (**A**) IAV derived from MDCK cells (red circles) blunts Spn growth compared to Spn grown in THY without treatment (black squares), and this effect is independent of cell culture media used in MDCK culture (blue triangles) Data represent n=8 per condition analyzed by two-way ANOVA, for clarity only significance at 3 h is shown (****p<0.0001). (**B**) MS analysis of most abundant (top 15) non-bacterial proteins present in the experimental media THY containing influenza A virus.

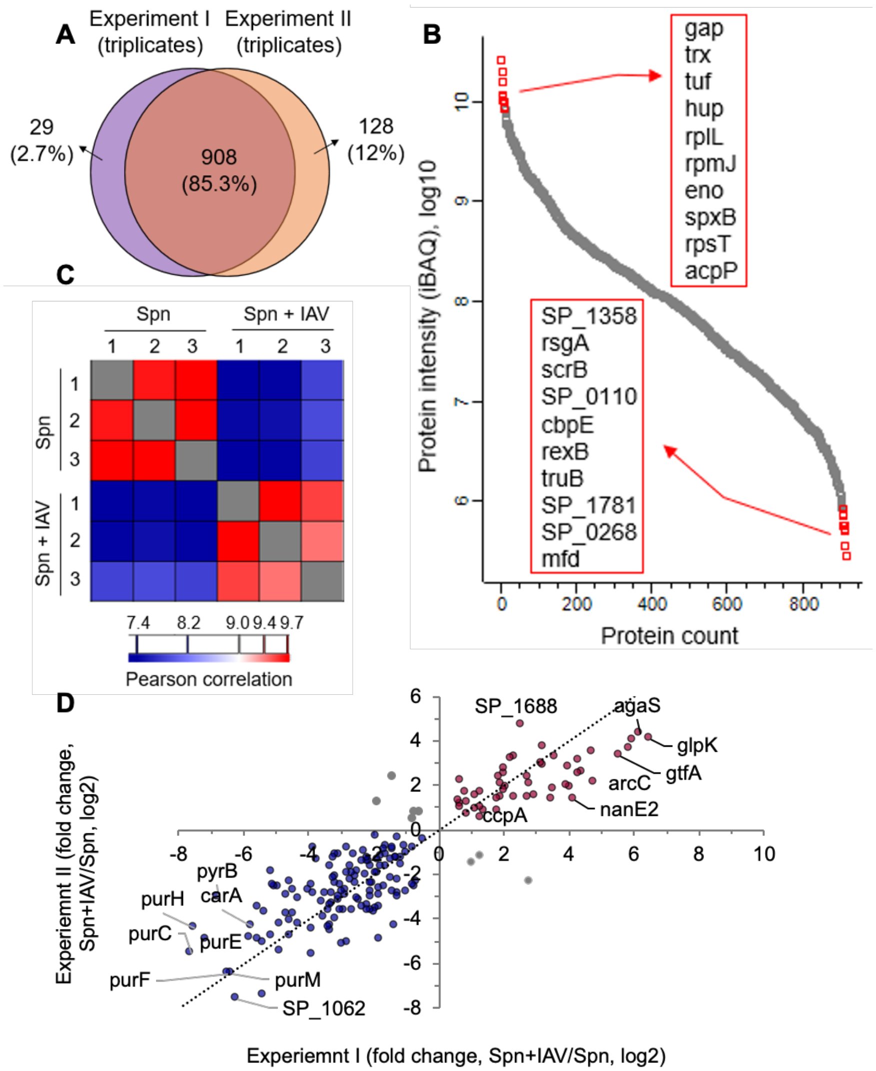


**Figure S4. Quantitative evaluation of *Spn* proteomics.** (**A**) Overlaps between two independent experiments. Each contained three biological replicates. (**B**) Dynamic range of the quantified Spn proteome. The tope 10 most and least abundant proteins were highlighted in the plot. (**C**) Pearson correlation among the replicates and the groups. (**D**) Correlation plot of significantly regulated proteins ($p < 0.05$) from the two experiments.

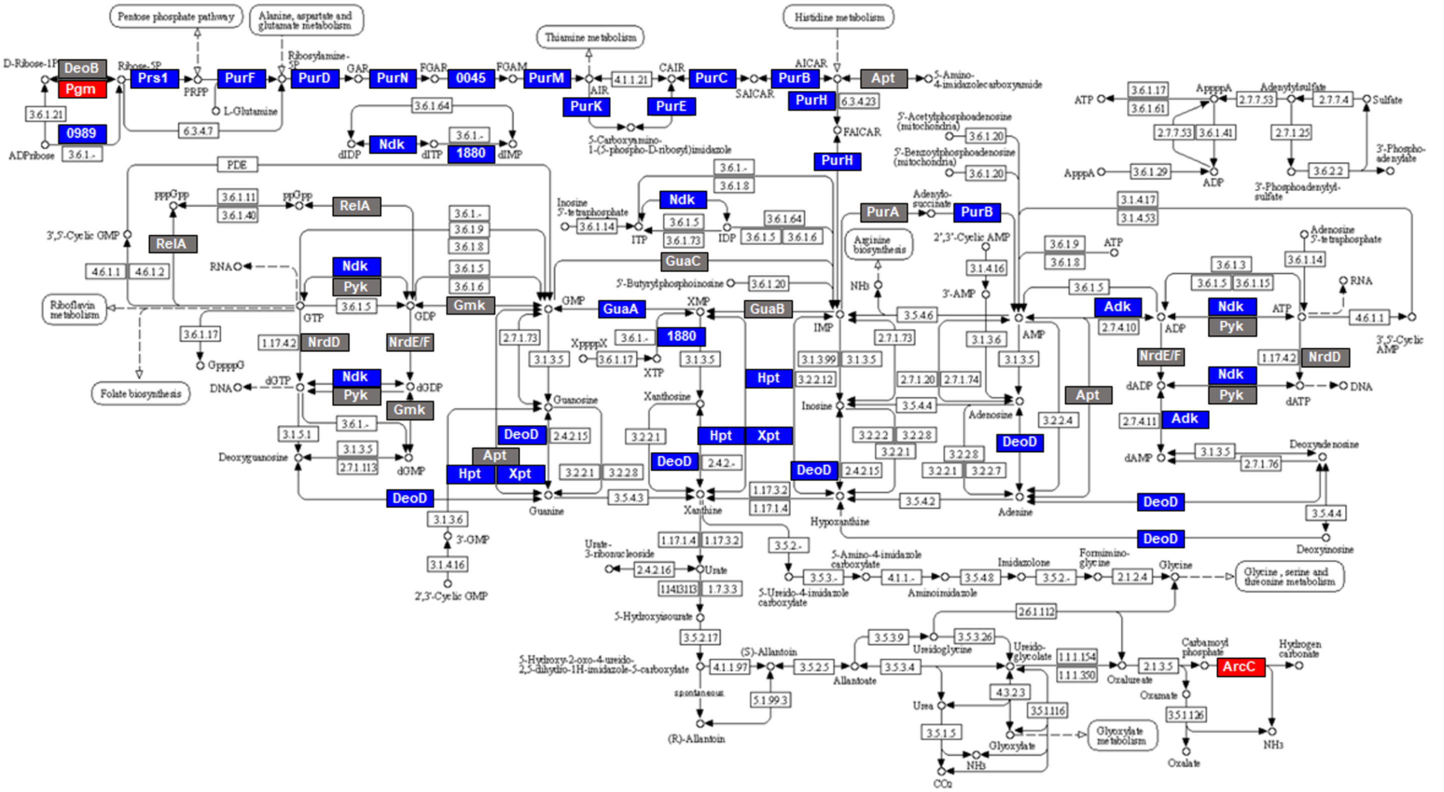


**Figure S5. Purine metabolism pathway.** *Spn* proteins participate in the pathway are in colored box. Red and blue indicates up- or down-regulation when *Spn* co-incubates with IAV, respectively. Gray indicates no significant changes or not identified in the proteome.

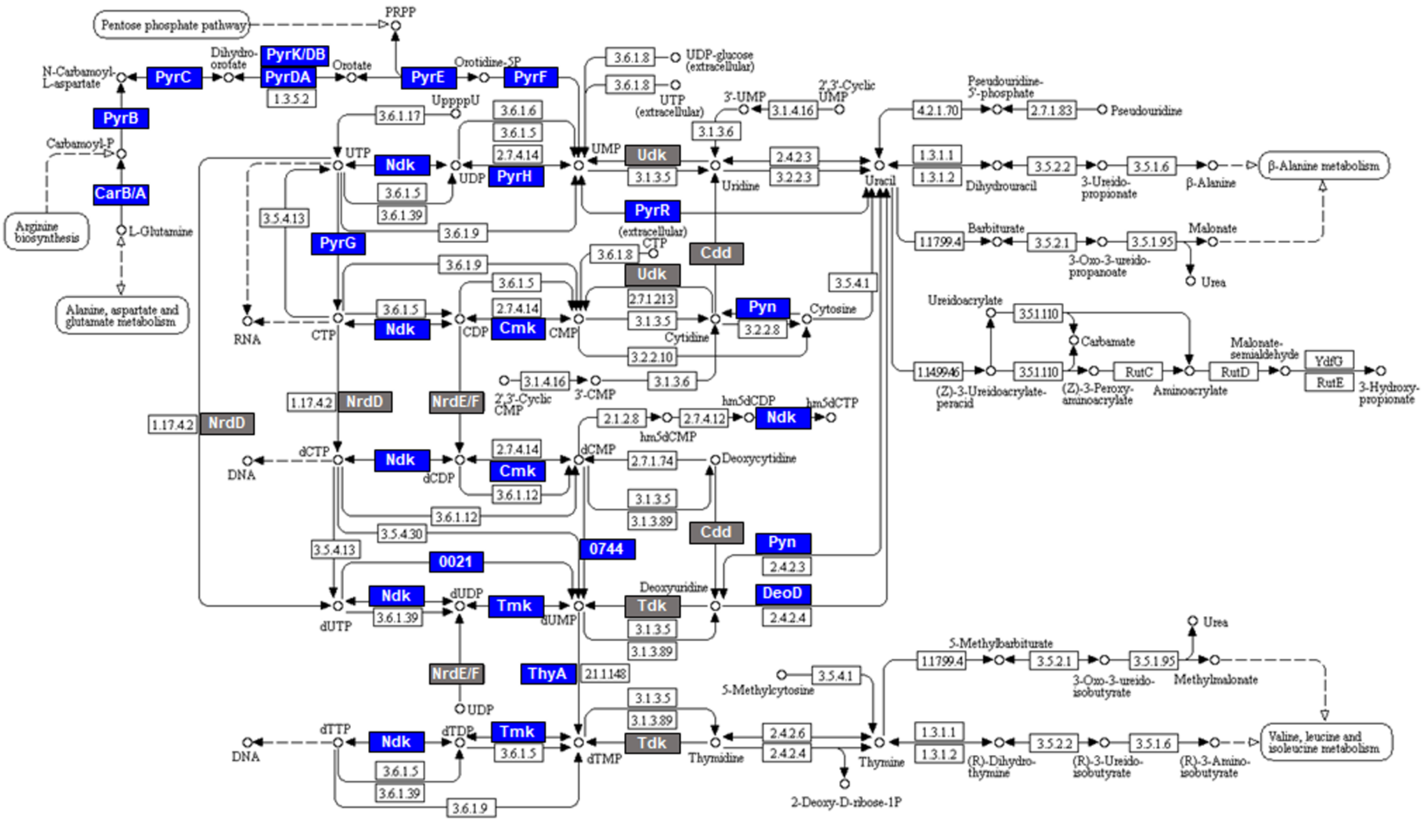


**Figure S6. Pyrimidine metabolism pathway.** *Spn* proteins participate in the pathway are in colored box. Red and blue indicates up- or down-regulation when *Spn* co-incubates with IAV, respectively. Gray indicates no significant changes or not identified in the proteome.

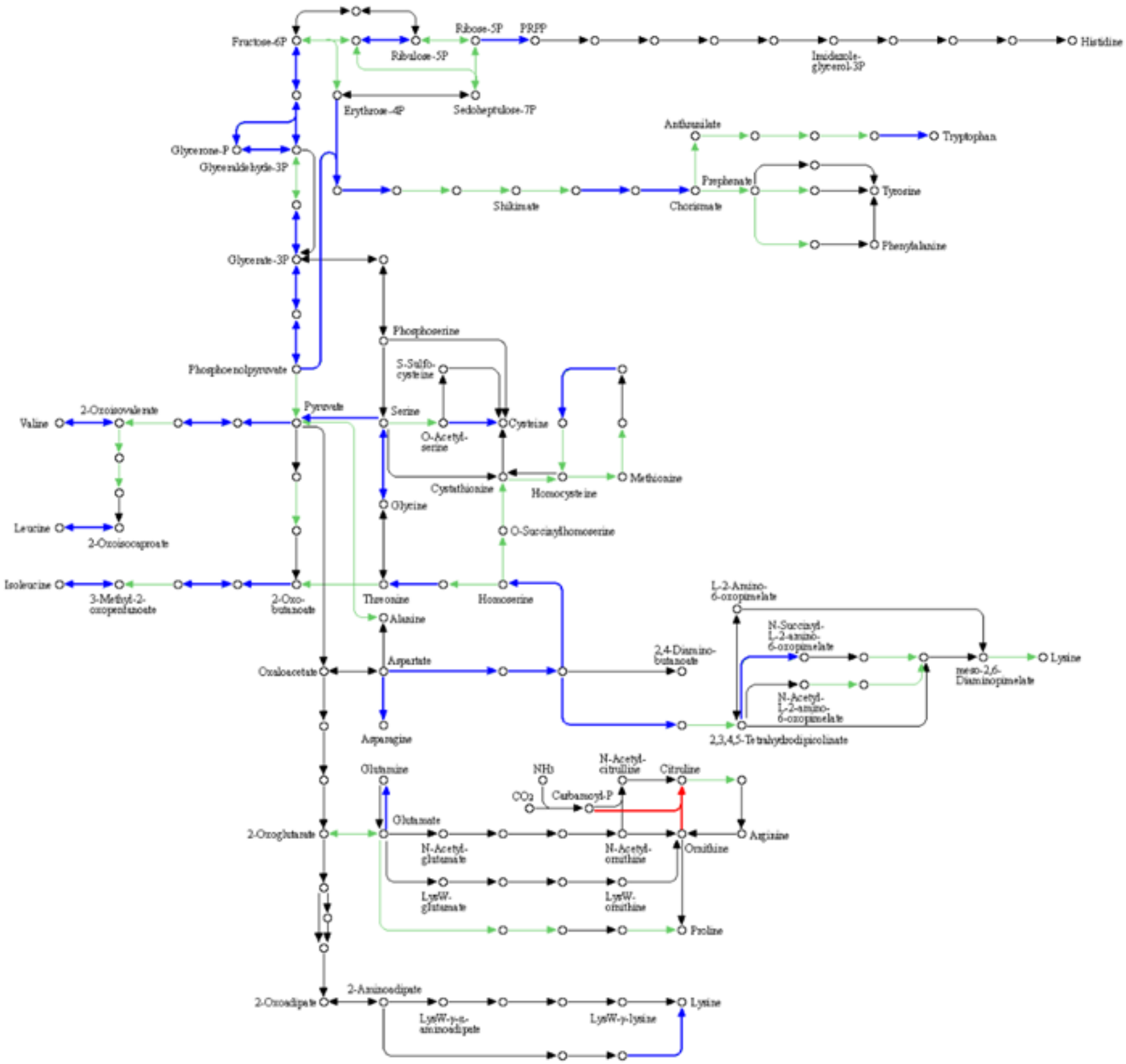

**Figure S7. Amino acids biosynthesis pathway.** *Spn* proteins participate in the pathway are in colored box. Red and blue indicates up- or down-regulation of *Spn* proteins co-incubated with IAV, respectively.

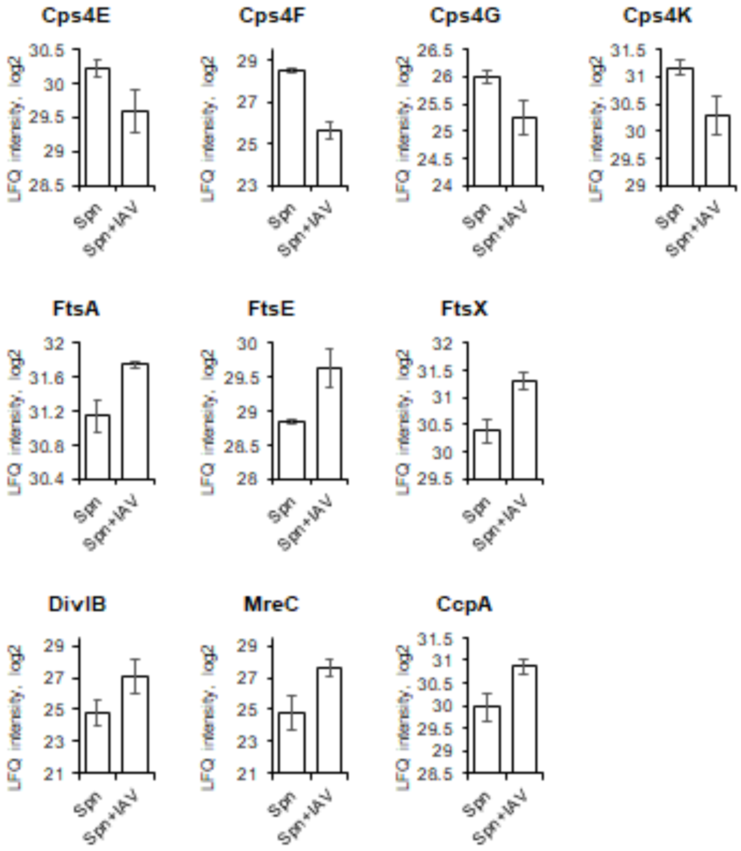


**Figure S8. Proteins involved in capsule synthesis, cell division, and carbohydrate metabolism were differentially regulated in *Spn* upon IAV co-incubation.** Representative significantly changed *Spn* proteins were plotted with their protein intensity (quantified by LFQ, log2-transformed) in the presence or absence of IAV.

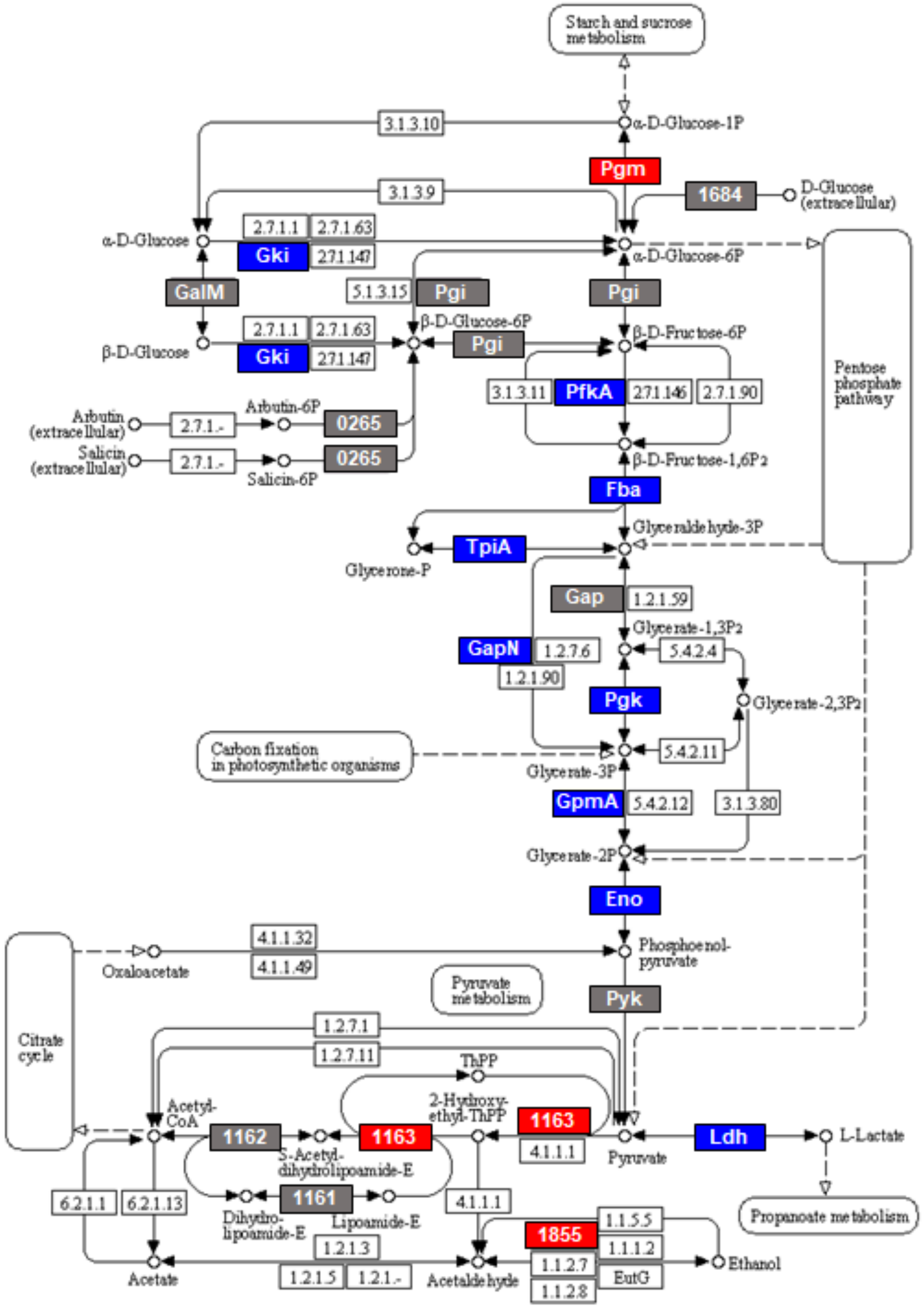


**Figure S9. Glycolysis/gluconeogenesis pathway.** *Spn* proteins participate in the pathway are in colored box. Red and blue indicates up- or down-regulation when *Spn* co-incubates with IAV. Gray indicates no significant changes or not identified in the proteome.

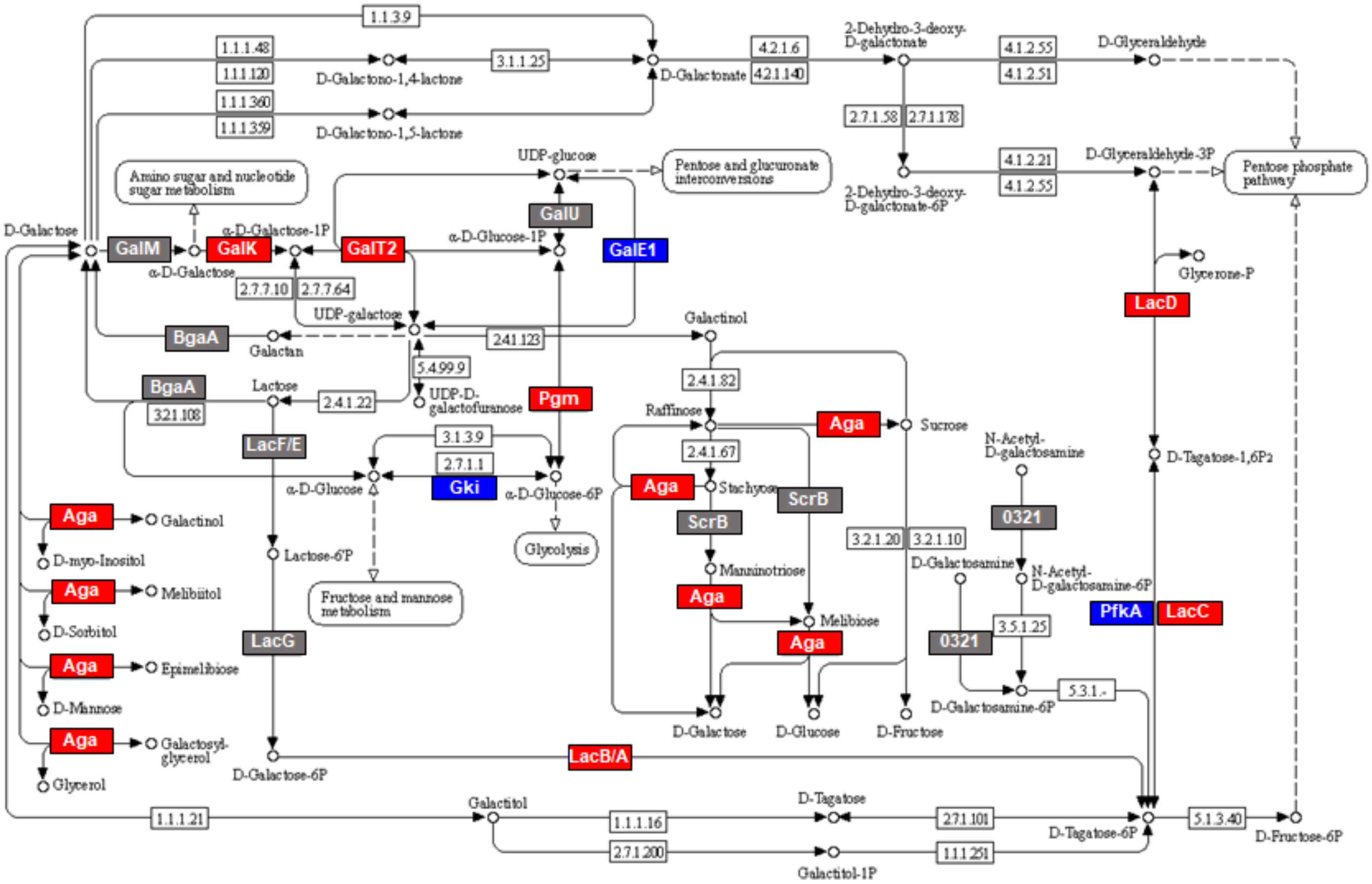


**Figure S10. Galactose metabolism pathway.** *Spn* proteins participate in the pathway are in colored box. Red and blue indicates up- or down-regulation when *Spn* co-incubates with IAV, respectively. Gray indicates no significant changes or not identified in the proteome.

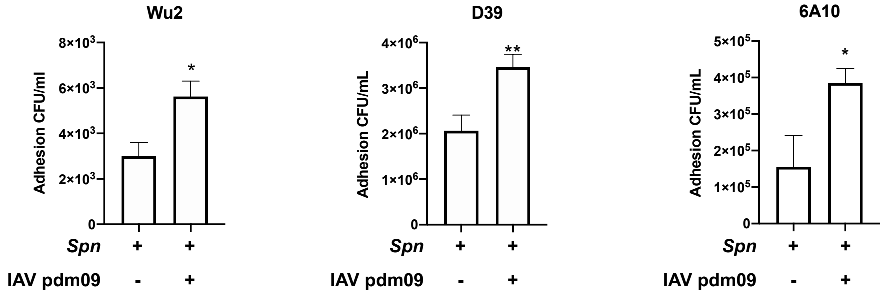


**Figure S11: IAV directly increases on *Spn* adhesion independent of serotype.** Adhesion assay of *Spn serotype* WU2, 6A10 and D39 to A549 cells was increased upon incubation with IAV (H1N1 A/California/7/2009 [pdm09]) for 1 hour prior to challenge of cells. Kruskal-Wallis test with Dunn's multiple comparison post-test. Asterisks denote the level of significance observed: * = p ≤ 0.05; ** = p ≤ 0.01.
