## Supplementary material for "A multiomics analysis of direct interkingdom dynamics between influenza A virus and *Streptococcus pneumoniae* uncovers host-independent changes to bacterial virulence fitness": Table 2

**Name	Sequence**

SP16s-1R	/5Biosg/GCGTTCTACTTGCATGTATTAGGCACGCCGCCAGCGTTCG

SP16s-2R	/5Biosg/TCCATTGCCGAAGATTCCCTACTGCTGCCTCCCGT

SP16s-3R	/5Biosg/ACCGCGGCTGCTGGCACGTAGT

SP16s-4R	/5Biosg/ACAGCGTGGACTACCAGGGTAT

SP16s-5R	/5Biosg/ACCACATGCTCCACCGCTTGTGCGGGCCCCCG

SP16s-6R	/5Biosg/ACCCAACATCTCACGACACGAGCTGACGACA

SP16s-7R	/5Biosg/GGGCGGTGTGTACAAGGCCCGGGA

SP16s-8R	/5Biosg/TTAAGAGATTAGCTTGCCGTCACCGGCTTGC

SP23s-1R	/5Biosg/ACCTTTCCCTCACGGTACTGGTTCACTATCGGTCA

SP23s-2R	/5Biosg/ACTCGCCGGTTCATTCTACAAAAGGCACGCTCTCACC

SP23s-3R	/5Biosg/TCGGAGAGAACCAGCTATCTCCAAGTTCGTTTGGA

SP23s-4R	/5Biosg/ATAGCTGCTTCTAAGCTAACATCCTA

SP23s-5R	/5Biosg/TAGTACAGGAATATCAACCTGTTGTCCATCGGATACACC

SP23s-6R	/5Biosg/TACCTGTGTCGGTTTGCGGTACGGG

SP23s-7R	/5Biosg/TCGTGCGGGTCGGAACTTACCCGACAAG

SP23s-8R	/5Biosg/GAGCCGACATCGAGGTGCCAAACC

SP23s-9R	/5Biosg/CGACGGATAGGGACCGAACTGTCTCACGAC

SP23s-10R	/5Biosg/GTGCCAAGGCATCCACCGTGCGCCCT

SP23s-11R	/5Biosg/CCACTTCTAACCTATCTACCTGATCATCTCTCAGG

SP5s-1R	/5Biosg/GGGTACAGGTGTATCTCCTAGGCTATCGTCAC

SP5S-2R	/5Biosg/CTAAGCGACTTCCCTATCTCACAGGGGG

**Table S2. *rRNA Depletion Primers*.**
