## Supplementary material for "A multiomics analysis of direct interkingdom dynamics between influenza A virus and *Streptococcus pneumoniae* uncovers host-independent changes to bacterial virulence fitness": Table 1

| **Gene** | **Protein description** | **Accession** | **Phosphosite Upregulated, Downregulated, *un-reported** |
| --- | --- | --- | --- |
| Cps4E | Capsular polysaccharide biosynthesis protein | A0A0H2UNI5 | Y89 |
| CpsD | Tyrosine-protein kinase | Q9AHD2 | Y221, S220, Y215, Y224, Y218 |
| DivIVA | Cell division protein | A0A0H2UR19 | **S224**, ***S205, *T201** |
| Fba | Fructose-bisphosphate aldolase | P0A4S1 | **S211** |
| FtsA | Cell division protein | A0A0H2UR24 | T404 |
| FtsZ | Cell division protein | A0A0H2UR32 | **T7**, ***S391, *T356** |
| Gap | Glyceraldehyde-3-phosphate dehydrogenase | A0A0H2US80 | T211 |
| GpmA | 2,3-bisphosphoglycerate-dependent phosphoglycerate mutase | P0A3Y3 | S144 |
| GpsB | Cell cycle protein | Q97SI9 | **S107**, ***T79** |
| MapZ | Mid-cell-anchored protein Z | A0A0H2UNG3 | **T27**, ***T78, *S80, *T67** |
| MltG | Endolytic murein transglycosylase | A0A0H2UQS8 | **T137**, **T138**, T157, T155, Y136, Y125 |
| Ndk | Nucleoside diphosphate kinase | P65536 | **S123** |
| NrdE | Ribonucleoside-diphosphate reductase | A0A0H2UQ44 | T155 |
| PtsH | Phosphocarrier protein HPr | A0A0H2UQ34 | **T12,** T20 |
| RpmC | 50S ribosomal protein L29 | P0A483 | T47 |
| SP_0095 | UPF0176 protein SP_0095 | SP_0095 | T116 |
| SP_0192 | UPF0297 protein SP_0192 | SP_0192 | T7 |
| SP_0571 | Cell filamentation protein Fic-related protein | SP_0571 | Y38, Y14 |
| SP_0877 | PTS system, fructose specific IIABC components | A0A0H2UPH7 | S207, T55 |
| SP_0990 | Uncharacterized protein | A0A0H2UPX4 | **T18**, ***T43, *T67** |
| SP_1110 | Riboflavin biosynthesis protein | A0A0H2UPY5 | T246 |
| SP_1339 | Uncharacterized protein | A0A0H2UQE9 | S30 |
| SP_1368 | Psr protein | A0A0H2UQH2 | Y24 |
| SP_1804 | Putative general stress protein 24 | A0A0H2URD5 | S195, S 166, S170 |
| SP_2040 | Putative jag protein | A0A0H2USA3 | T89 |
| Ssb | Single-stranded DNA-binding protein | P66854 | T144 |
| StkP | Serine/threonine-protein kinase | Q97PA9 | **T305**, **T303**, **T330**, ***T293, *T295** |
| TufA | Elongation factor Tu | P64030 | T387, **S52**, **S43** |

**Table S1. *Spn* phosphoproteins identified in this study.** *Spn* strain TIGR4 phosphoproteins, their Uniprot accession numbers and phosphosites identified in this study are listed. Phosphosites that are up- or down-regulated in the presence of IAV are shown in bold or underlined, respectively.
